# The *C. elegans* ephrin EFN-4 functions non-cell autonomously with heparan sulfate proteoglycans to promote axon outgrowth and branching

**DOI:** 10.1101/022756

**Authors:** Alicia A. Schwieterman, Alyse N. Steves, Vivian Yee, Cory J. Donelson, Aaron Pital, Taylor Voyles, Austin M. Howard, Danielle E. Ereddia, Kelsie S. Effrein, Jonathan L. McMurry, Brian D. Ackley, Andrew D. Chisholm, Martin L. Hudson

## Abstract

The Eph receptors and their cognate ephrin ligands play key roles in many aspects of nervous system development. These interactions typically occur within an individual tissue type, serving either to guide axons to their terminal targets or to define boundaries between the rhombomeres of the hindbrain. We have identified a novel role for the *Caenorhabditis elegans* ephrin EFN-4 in promoting primary neurite outgrowth in AIY interneurons and D-class motor neurons. Rescue experiments reveal that EFN-4 functions non-cell autonomously in the epidermis to promote primary neurite outgrowth. We also find that EFN-4 plays a role in promoting ectopic axon branching in a *C. elegans* model of X-linked Kallmann syndrome. In this context, EFN-4 functions non-cell autonomously in the body wall muscle, and in parallel with HS biosynthesis genes and HSPG core proteins, which function cell autonomously in the AIY neurons. This is the first report of an epidermal ephrin providing a developmental cue to the nervous system.

## INTRODUCTION

Accurate development of the central nervous system requires contributions from both the extra-cellular environment, in the form of guidepost cues and secreted guidance molecules, and from contact with adjacent neurons or other tissues, in the form of cell surface receptors that can detect and transduce navigation cues. Guideposts typically take the form of specific tissue types such as the ventral midline of *C. elegans* or *Drosophila*, which provides a permissive environment for neurons to migrate along, whilst secreted guidance molecules provide spatial information and navigation instructions in the form of repulsive or attractive cues (Chilton 2006; Killeen and Sybingco 2008). Many studies have identified how individual neurons and guidance cues act to provide spatial and navigation information, but how does a single cell that possesses multiple axon guidance receptors find its target when exposed to multiple guidance cues? The nervous system of the nematode *C. elegans* provides a simple and defined model to examine the interplay between multiple neuronal guidance systems. The morphology and connectivity of *C. elegans* neurons have been established by serial section electron microscopy and the *C. elegans* genome possesses orthologs of most vertebrate axon guidance and guidepost genes (White et al. 1986; Bargmann 1998; Chisholm and Jin 2005; Ackley 2014).

One of the most important classes of axon guidance molecules is the Eph receptor tyrosine kinases and their cognate ligands, the ephrins (Flanagan 2006; Lisabeth et al. 2013; Cayuso et al. 2014). Eph receptors and ephrins are required for the accurate connectivity of many parts of the vertebrate brain, and also have roles in cell adhesion and embryonic morphogenesis (Klein 2012; George et al. 1998; Chin-Sang et al. 1999; Chin-Sang et al. 2002). We previously showed that the *C. elegans* ephrin EFN-4 functions in concert with the KAL-1/anosmin - heparan sulfate proteoglycan (HSPG) pathway to regulate neuroblast migration during embryonic development (Hudson et al. 2006). Anosmin is an extracellular matrix molecule that is highly conserved between humans and *C. elegans* (Rugarli et al. 2002; Bulow et al. 2002; Hu et al. 2003). Mutations in the human KAL1/anosmin gene lead to X-linked Kallmann syndrome, which is characterized by loss of sense of smell and the failure to undergo spontaneous puberty (Kallmann et al. 1944; Franco et al. 1991; Legouis et al. 1991; Dode and Hardelin 2009). Curiously, rodents appear to lack orthologs of KAL1, making *C. elegans* one of the few model systems with which to investigate this disease. Over-expression of KAL-1 in the *C. elegans* central nervous system creates a highly penetrant, cell autonomous ectopic branching phenotype (Bulow et al. 2002). This phenotype is strongly suppressed by mutations in HSPG biosynthetic enzymes and in vitro binding studies have confirmed that KAL-1 can bind the HSPGs *sdn-1*/syndecan and *gpn-1*/glypican (Bulow et al. 2002, Hudson et al. 2006; Tornberg et al. 2012). HSPGs are required for many aspects of nervous system development in both vertebrates and invertebrates, including cell migration, axon guidance and synaptogenesis (Rhiner et al. 2005; Van Vactor et al. 2006; Kinnunen 2014). Considering the importance of both HSPGs and the Eph/ephrin signaling during nervous system development, surprisingly little research has been dedicated to possible interactions between these pathways (Irie et al. 2008; Holen et al. 2011). In this study, we focus on the interplay between ephrins and HSPGs in the development of *C. elegans* AIY interneurons. We show that the ephrin EFN-4 is required non-cell autonomously to promote AIY primary neurite outgrowth, functioning in parallel with SDN-1/syndecan in this process. We also show that in a *C. elegans* model of X-linked KS, EFN-4 plays a role in promoting AIY ectopic neurite branching, where it again functions non-cell autonomously. Finally, we show that SDN-1/syndecan and GPN-1/glypican have cell autonomous yet mutually antagonistic roles in ectopic neurite formation. This is the first report of an ephrin acting non-cell autonomously from the epidermis to regulate neurite outgrowth and branching.

## MATERIALS AND METHODS

### Strains and maintenance

*C. elegans* strains were grown on nematode growth medium plates (NGM Lite) at 20°C according to Brenner (1974). All analyses were conducted at 20°C unless otherwise noted. The following mutations were used in the course of this work:

LGI: *kal-1(gb503); mab-20(ev574);* LGII: *vab-1(e2027);* LGIII: *hse-5(tm472);* LGIV: *efn-4(bx80, e36, e660, e1746ts), lad-2(tm3056);* LGX: *sul-1(gk151), hst-2(ok595), hst-6(ok273), sdn-1(ok449), sdn-1(zh20), gpn-1(ok377), lon-2(e678).*

The following transgenes were used in the course of this work:

LGII: *juIs76[P-unc-25-GFP + lin-15(+)],* LGIV: *mgIs18[P-ttx-3-GFP], otIs76[P-ttx-3-kal-1(+)*+ *P-unc-122-GFP], juIs109[P-efn-4-efn-4::GFP + lin-15(+)], oxTi420[P-eft-3-mCherry::tbb-2-*3’UTR + *Cbr-unc-119(+)],* LGX: *oxIs12[P-unc-47-GFP lin-15(+)]*, *otEx331[P-lad-2-GFP* +*pha-1(+)]*

The following extrachromosomal arrays were generated in the course of this work:

*kenEx4, kenEx5, kenEx6 [P-myo-3-efn-4 + P-myo-3-mCherry]; kenEx9, kenEx10, kenEx11 [p-myo-2-efn-4 + P-myo-2::mCherry]; kenEx12, kenEx13, kenEx14 [efn-4 genomic + P-ttx-3-RFP]; kenEx16, kenEx17, kenEx18 [P-ttx-3-lon-2 + P-ttx-3-RFP]; kenEx19, kenEx20 [P-unc-119-efn-4 + P-ttx-3::RFP]; kenEx21, kenEx23 [P-elt-3-efn-4 + sur-5::GFP + P-ttx-3-RFP]; kenEx27, kenEx28, kenEx29 [P-elt-3-lon-2 cDNA + P-ttx-3-RFP]; juEx1335, juEx1336, juEx1337 [P-ttx-3-GFP::sdn-1 cDNA + P-ttx-3-RFP]; juEx1338, juEx1339, juEx1340 [P-gpn-1-GFP::gpn-1 cDNA + P-ttx-3-RFP]; juEx1341, juEx1342, juEx1343[P-AIY-efn-4 cDNA + P-ttx-3-RFP]*; *juEx1358, juEx1359, juEx1360 [P-ttx-3-gpn-1 cDNA + P-ttx-3-RFP]; lhEx174, lhEx175, lhEx176 [P-lon-2-lon-2 cDNA + P-ttx-3-RFP].*

### Neuroanatomy

Neuronal morphology was scored using GFP or RFP expression from reporter genes. Animals were either imaged directly using a Zeiss Axioskop compound microscope, or off-line, from z-stacks of images acquired on a Zeiss LSM700 confocal microscope. Only axon branches exceeding 5 μm in length were included for analysis (Bulow et al. 2002). Distance was measured with Zeiss ZEN Blue software tools or the Segmented Line function of ImageJ (Rasband, 1997-2014). Primary neurite outgrowth defects were scored as “shortstop” if they failed to reach their terminal target on the dorsal side of the nerve ring as defined by a visible gap of greater than 3 microns.

### Transgenic rescue of efn-4 and HSPG phenotypes

To assay for rescue of AIY phenotypes, transgenic lines were generated bearing either genomic clones or tissue-specific rescue plasmids in conjunction with a co-injection marker. All plasmids were generated in this work unless otherwise noted: pCZ148 (*efn-4* genomic clone, Chin-Sang et al. 2002), pMH340-3 (*P-ttx-3-efn-4* cDNA), pMH838 (*P-unc-119-efn-4* cDNA), pMH853 (*P-myo-2-efn-4* cDNA), pLC681 (*P-myo-3-efn-4* genomic, kind gift of Lihsia Chen), pMH833 (*P-elt-3-efn-4* cDNA), pMH306-7 (*P-ttx-3-GFP::sdn-1* cDNA), pMH265 (*P-gpn-1-GFP::gpn-1 cDNA*), pMH274 (*P-ttx-3-gpn-1* cDNA), pHW474 (*P-lon-2-lon-2* cDNA, Gumienny et al. 2007), pTG102 (*P-elt-3-lon-2* cDNA, Gumienny et al. 2007), pMH644 (*P-ttx-3-lon-2* cDNA). *efn-4* plasmids were injected at 1 or 5 ng.μl^-1^ into *efn-4(bx80) mgIs18 otIs76*. HSPG plasmids were injected into *hspg(null) mgIs18 otIs76* at 1 or 5 ng.μl^-1^.The following co-injection plasmids were used: pTTX-3-RFP (AIY neuron RFP, Hobert et al. 1997), pTG96 (all nuclei GFP, Yochem et al. 1998), pPD131-68 (body wall muscle mCherry, kind gift of A. Fire), pPD95-91 (pan-neuronal GFP, Fire et al. 1990), pBA183 (pharyngeal mCherry). Plasmid construction details and sequences are available on request. See Supplemental data S1 and S2 for details of *efn-4* and HSPG transgenic line construction.

### Statistical analysis

Statistical significance was determined using the Z-test with an alpha level of 0.05. Differences were considered significant if P < 0.05 (*), p < 0.01 (**) or p < 0.001 (***). Error bars on graphs show the standard error of proportion. Bonferroni corrections were not applied as either multiple alleles or multiple transgenic lines were assayed in parallel.

### Expression plasmid construction

The following plasmids were used to express and secrete proteins from 293T cells: pCZ144 (VAB-1 extra-cellular domain::AP Chin-Sang et al. 1999), pMH180 (Fc::mycHis), pMH873 (EFN-1::Fc::mycHis), pMH874 (EFN-4::Fc::mycHis). See supplementary data S3 for details on plasmid construction. Plasmid sequences are available on request.

### Cell Culture

HEK293T cells were maintained in Dulbecco’s Modified Eagle’s Medium (DMEM) (Caisson Laboratories Inc.) supplemented with 10% ultra-low IgG Fetal Bovine Serum (Life Technologies), 1mM L-Glutamine (Caisson Laboratories Inc.), and 1x Penicillin-Streptomycin (Caisson Laboratories Inc.).

### Transient Transfection of EFN-1, EFN-4 and VAB-1 expression plasmids

Expression plasmids were transfected into HEK293T cells using Lipofectamine® 2000 Transfection Reagent (Life Technologies) and Opti-MEM® Reduced Serum Medium (Life Technologies) according to the manufacturer’s protocol. Final DNA concentrations were 10ng/μl. Transfections were incubated at 37°C in 5% CO_2_ atmosphere for 48 hours. Supernatants were harvested, centrifuged to remove cellular debris, then either used directly or concentrated using a 10kDa cut off centrifugal filter (Centricon). Protein production was confirmed by either Western blotting to detect myc-tags of the transfected ephrin-Fc proteins, or by AP-turnover assay for VAB-1(ECD)::AP.

### Biolayer Interferometry

Biolayer interferometry measurements were made on a FortéBio Octet -QK instrument using anti-human IgG Fc capture (AHC) biosensors (FortéBio, Inc.). Assays were performed in 96-well micro-plates at 25°C. All volumes were 200 μl. Biosensors were conditioned for 3 minutes in Opti-MEM media prior to loading. Ephrin-Fc ligands or Fc controls were bound onto AHC biosensors for 60 minutes then a baseline established in Opti-MEM for 15 minutes. Sensors were then incubated in VAB-1::AP tissue culture supernatant for 60 minutes (analyte association phase), followed by incubation in Opti-MEM for 60 minutes (dissociation phase).

## RESULTS

Our previous work showed that *efn-4* mutations synergize with *kal-1* in neuroblast migrations during embryonic morphogenesis, suggesting EFN-4 and KAL-1 play closely related roles in cell migration or adhesion (Hudson et al. 2006). To learn whether EFN-4 and KAL-1 also function together in neural development we tested whether *efn-4* mutations displayed genetic interactions with *kal-1(lf)* and *kal-1(gf)* mutations during axon outgrowth and branching.

### EFN-4 is required for neural development

The AIY neurons are a bilaterally symmetric pair of interneurons that are born about 365 minutes post-fertilization (Bao et al. 2006; Murray et al. 2008). We used *mgIs18*, an AIY-specific GFP-transgene driven from a *ttx-3-*intronic enhancer element, to monitor development of the AIY interneurons in embryos, larvae and adults (Altun-Gultekin et al. 2001). Following ventral enclosure, the AIY cell bodies lie adjacent to myoblasts, the developing pharynx and also the ventral epidermis. They begin to extend their primary neurites anteriorly during the lima bean stage of embryonic development then dorsally at the 1.5 fold stage (Christensen et al. 2011 and M. L. Hudson, unpublished observations). The AIYL and R cell bodies ultimately lie in the ventral ganglion beneath the posterior pharyngeal bulb. Their main axons, which we refer to in this study as the primary neurites, converge anteriorly at a plexus-like area where they enter the nerve ring. Within the plexus the AIYs are flattened immediately adjacent to the cephalic sheath cells, where they occasionally extend very short collateral branches. After entering the nerve ring the AIY primary neurites extend circumferentially to the dorsal midline where they contact one another via a gap junction (Figure 1a, b; White et al. 1986).

**Figure 1.**
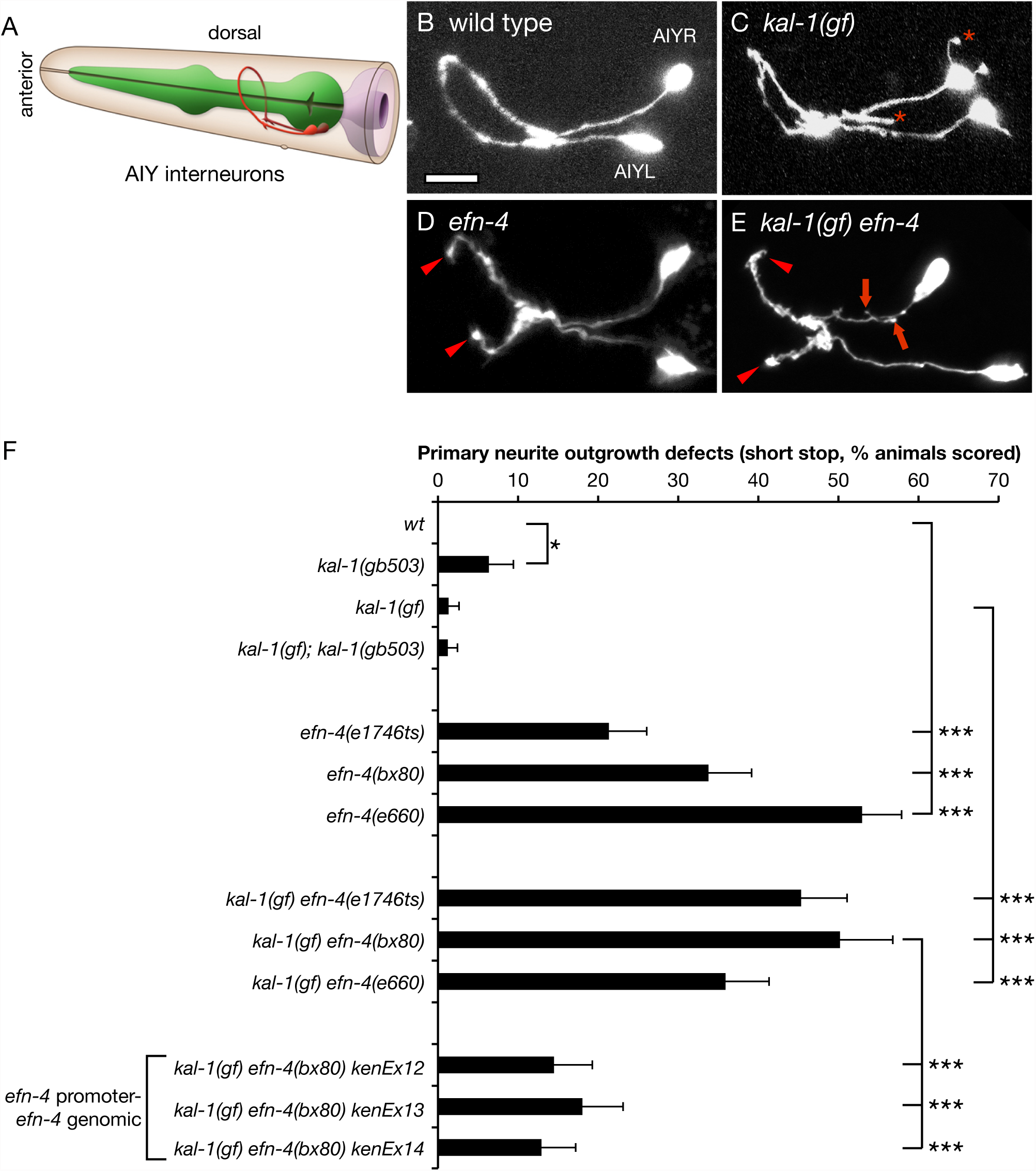
*efn-4* mutations have defects in primary neurite outgrowth and suppress KAL-1-dependent axon branching. (a) Cartoon depiction of AIY interneuron morphology in relation to the pharynx and epidermis. AIY neurons are bilaterally symmetrical unipolar cells that extend a single axon anteriorly. The axons contact their contralateral partner on the ventral side of the nerve ring beneath the pharynx before extending anteriorly, where they again make contact on the dorsal side of the worm (image adapted from www.wormatlas.org). (b) Confocal micrograph of AIY neuron morphology in a wild type background. (c) *kal-1(gf)* worms over-express KAL-1 in the AIY neurons, causing a highly penetrant ectopic axon branching phenotype (asterisks). (d) AIY neuron morphology in an *efn-4(bx80)* mutant background. Arrowheads show premature axon outgrowth termination, resulting in a failure to make contact on the dorsal side of the nerve ring. (e) *kal-1(gf)* axon branching is suppressed in *efn-4* mutants. Arrows show short, stubby axon branches. Arrowheads show premature axon termini. Scale bar = 10mm. (f) Graph of AIY axon outgrowth defects in *efn-4* mutant backgrounds. All alleles tested show axon outgrowth defects and are generally more penetrant in a *kal-1(gf)* background. *kenEx12 – 14* are transgenic rescue arrays containing an *efn-4* genomic clone that includes 5kb of the *efn-4* promoter region. Statistical significance was determined using the z-test. * p<0.05; *** p < 0.001. Error bars are standard error of proportion.

We find that all *efn-4* loss of function alleles show strong premature axon termination defects in the AIY interneurons, which manifest as failure of the primary neurites to meet at the dorsal midline (Figure 1d, f). We refer to this as a “shortstop” phenotype, as the overall trajectory of the axons is correct, but the axons stop short of their correct termination site. The phenotypic penetrance roughly correlates with the allelic strength described by Chin-Sang et al. (2002) with *bx80* and *e660* showing 34% and 53% shortstop phenotype respectively, whereas the *e1746ts* allele shows only 21% short stop. This is in contrast to *mgIs18* controls that show no shortstop phenotype (90 animals scored). We also observed that AIY axon outgrowth defects in *efn-4* mutants was intrinsically temperature sensitive, irrespective of the allele tested (Table S4). For instance, *efn-4(bx80)* showed only 23% outgrowth defects at 15°C but increased to 34% and 32% at 20°C and 25°C respectively. For consistency, all remaining data were scored at 20°C.

We also assayed *efn-4* shortstop phenotypes in a *kal-1(gf)* background (Figure 1e, f). Again, all three *efn-4* alleles showed highly penetrant shortstop phenotypes, although *efn-4(e1746ts)* mutants displayed significantly stronger phenotypes in a *kal-1(gf)* background, whereas *efn-4(e660)* were significantly weaker. These differences may reflect the overall sensitivity of this phenotype to temperature variation. Considering that all three *efn-4* alleles examined had strong shortstop phenotypes irrespective of the *kal-1(gf)* background, and that *kal-1(gf)* alone has no phenotype, we conclude that KAL-1 over-expression has no overt influence on primary neurite outgrowth.

To confirm that EFN-4 is required for AIY primary neurite outgrowth, we used germ line microinjection to reintroduce an *efn-4* genomic clone into *efn-4(bx80)* null mutants. This rescued the AIY shortstop phenotype from 50% (27/54 animals tested) down to around 15% depending on the transgenic line in question (Figure 1f, 3 independent lines, p < 0.001). These rescue support our genetic loss of function analysis, further confirming that EFN-4 is required for AIY primary neurite outgrowth.

### EFN-4 promotes D-class motorneuron outgrowth but not axon guidance

EFN-4 is expressed in multiple tissues at various stages of the *C. elegans* life cycle and as such, could be required for the development of many neuron types including the ventral nerve cord (Chin-Sang et al. 2002). We used the *P-unc-25-GFP* reporter, *juIs76*, to investigate whether EFN-4 has roles in shaping the morphology of D-type motorneurons (Zhen and Jin, 1999). The 6 DD neurons migrate to the ventral midline during mid-embryogenesis and extend an axon anteriorly, which then branches midway along the axon, extends a commissural process circumferentially around the worm until it meets the dorsal midline where it again branches, extending processes to both the anterior and posterior (Figure 2a). DD neuron commissural migration is stereotyped in that the DD1 neuron extends its process to the left-hand side of the animal, whereas DD2-6 extends commissures to the right hand side. The 13 VD neurons are born late in the L1 larval stage and share similar morphology and guidance choices to the DD neurons; VD1 extends a process to the left whereas VD2-13 extends processes to the right (White et al. 1976). In both DD and VD neurons, the anterior and posterior processes of each cell make a gap junction contact with their respective anterior and posterior DD/VD neighbors and provide GABAergic inhibitory innervation to the body wall muscles to promote smooth sinusoidal locomotion (White et al. 1976). In wild type animals, the ventral and dorsal nerve cords appear as contiguous structures, with only 4% of animals (2/44) showing gaps in the dorsal D-neuron cord (Figure 2b). In *efn-4(bx80)*, 29% of animals (15/53) show at least one gap in dorsal cord morphology (Figure 2c, d). In addition, 8% of animals (4/53) show a “crooked” dorsal cord phenotype (Figure 2c) as opposed to the linear dorsal cord seen in wild type animals. We also examined left/right outgrowth choice of the commissural processes. In wild type, 7% of worms (3/47) show at least one axon where the commissural process navigates dorsally via the wrong side of the animal. In *efn-4* mutants, 9% of worms (5/58) show a commissural guidance defect (not significant). Taken together, we conclude that EFN-4 is required, in part, to promote the final stage of D-neuron outgrowth along the dorsal nerve cord, similar to its role in promoting the final stage of AIY axon outgrowth, but that it has no role in axon guidance. While it is possible that EFN-4 is required for the development of other neural cell types beyond the AIY interneurons and D-motorneurons, the remainder of this work will focus solely on EFN-4’s role in shaping AIY interneuron morphology.

**Figure 2.**
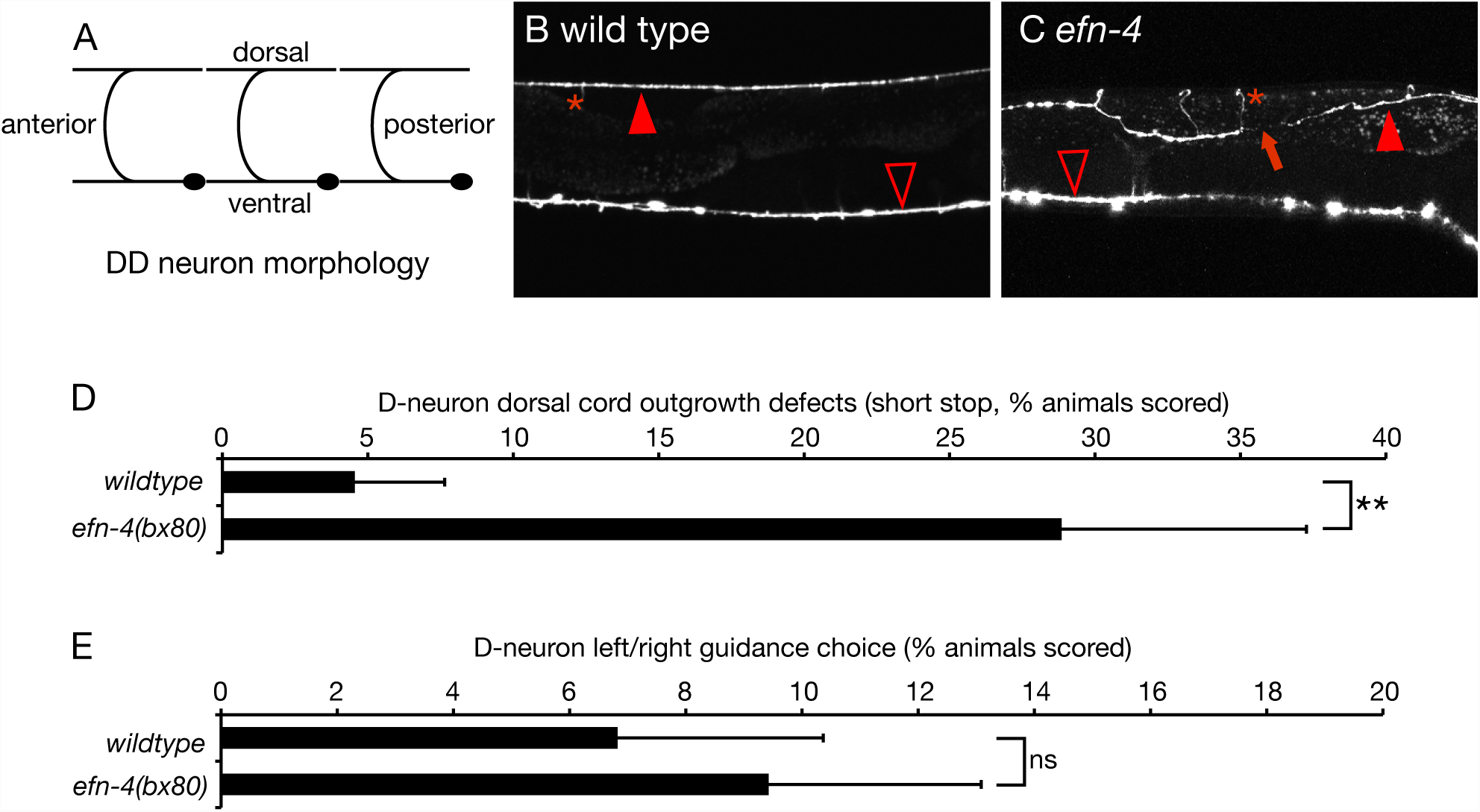
*efn-4* mutations show defects in D-neuron axon outgrowth. (a) Diagram of DD-neuron morphology (VD neurons are similar in shape but their commissures are located further away from the cell body). (b) Confocal micrograph of GFP expression from the D-neuron specific GFP marker *juIs76[P-unc25-GFP].* (c) *efn-4* mutants display defects in D-neuron outgrowth (arrow) and can also show twisted dorsal axon tracts. Filled arrows, dorsal nerve cord; open arrows, ventral nerve cord; asterisk, commissural processes. (d) Graph of D-neuron dorsal outgrowth defects and (e) left/right commissural guidance decisions in wild type versus *efn-4(bx80)* mutants. Statistical significance was determined using the z-test. ** p<0.01. Error bars are standard error of proportion.

### EFN-4 is required to promote kal-1(gf) ectopic branching

We previously showed that *efn-4* mutants exhibit strong genetic interactions with *kal-1*/anosmin mutants during embryonic ventral neuroblast migrations (Hudson et al. 2006). Over-expression of KAL-1 in the AIY neurons causes highly penetrant cell autonomous ectopic neurite branching (Bulow et al. 2002). The ectopic branches extend posteriorly from the ventral plexus of the AIYs, and may represent hypertrophied versions of the small branches normally formed in this region. We confirmed that *P-AIY-kal-1* (*otIs76*) causes ectopic AIY branch formation and also observed ectopic branches emanating from AIY cell bodies and occasionally the nerve ring process (Figure 1c and Figure 3). We also found that ectopic branch formation is temperature sensitive (Table S4 lists all strains and AIY phenotypes scored at different temperatures during the course of this work). At 20°C 55% of *mgIs18 otIs76* AIY neurons (84% of animals; n = 153) exhibited ectopic branches. We found that loss of function in *kal-1* suppresses this *kal-1(gf)* effect, supporting the interpretation that ectopic branches are due to excess of normal KAL-1 function. Interestingly, *kal-1(lf)* mutations themselves caused low penetrance branching of AIYs, suggesting that either reduction or elevation of KAL-1 function may affect branching.

**Figure 3.**
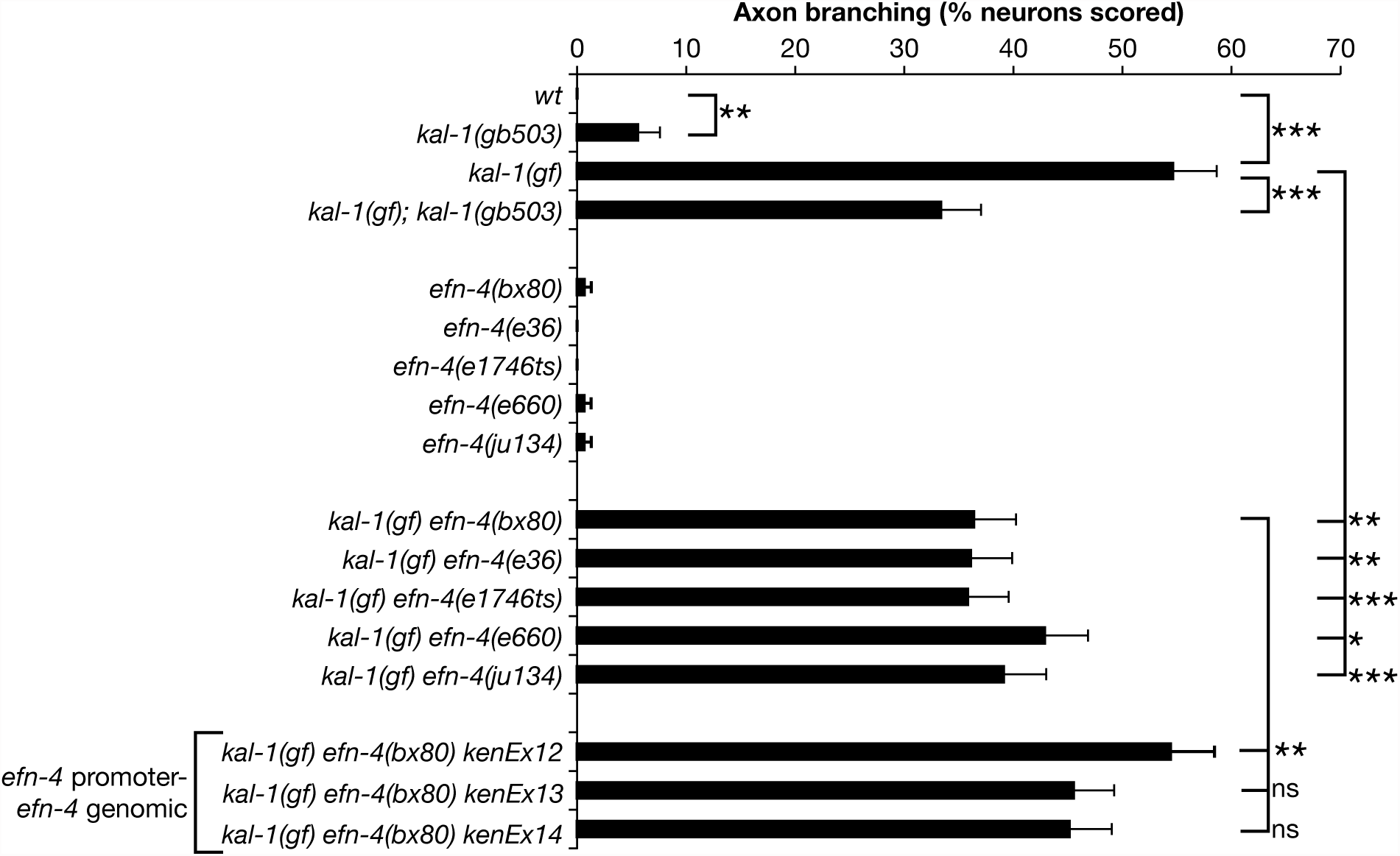
*efn-4* mutations suppress *kal-1(gf)-*induced axon branching. All *efn-4* alleles tested can strongly suppress *kal-1(gf)* axon branching relative to *kal-1(gf)* alone. *kenEx12 – 14* are transgenic rescue arrays containing an *efn-4* genomic clone that includes 5kb of the *efn-4* promoter region. Statistical significance was determined using the z-test. * p<0.05; *** p < 0.001. Error bars are standard error of proportion.

We next examined AIY ectopic branching in *kal-1(gf) efn-4(-)* lines. We found that all *efn-4* alleles significantly suppressed *kal-1(gf)* branching (Figure 1c, 1e and Figure 3). Reintroduction of an *efn-4* genomic clone into *kal-1(gf) efn-4(bx80)* mutants de-suppresses this phenotype, although the change was only significant in one of three lines tested (Figure 3). From this, we conclude that EFN-4 contributes to *kal-1(gf)* induced ectopic branching, but that other genes are likely to be required in this process.

To better understand what additional genes might contribute to AIY primary neurite extension and *kal-1gf)* induced ectopic branching, we examined AIY morphology in other genetic mutants known to affect axon outgrowth (Table 1). *plx-2*/plexin and *mab-20/*semaphorin have both been implicated in EFN-4 function during neuroblast migration, male tail sensory ray formation and axon guidance (Nakao et al. 2007; Wang et al. 2008). We found that *plx-2* mutants showed no defects in primary neurite outgrowth, nor did they affect *kal-1(gf)* ectopic axon branching. This was in sharp contrast to *mab-20* mutants, which showed highly penetrant neurite outgrowth defects, similar to *efn-4* mutants. This concurs with previous genetic studies, placing *efn-4* and *mab-20* in a common neurodevelopmental pathway (Chin-Sang et al. 2002; Ikegami et al. 2004; Nakao et al. 2007). The LAR-like receptor protein tyrosine phosphatase PTP-3 is required for embryonic morphogenesis, axon outgrowth and synapse formation (Harrington et al. 2002; Ackley et al. 2005). We found that *ptp-3(mu256)* mutants also exhibits defects in both primary neurite outgrowth and *kal-1(gf)* ectopic branching. We did not attempt further analysis of *efn-4* and *ptp-3* genetic interactions due to the strong synthetic-lethal phenotype observed between *ptp-3(mu256)* and all *efn-4* alleles tested (R. Harrington and A.D.C., unpublished observations). Mutations in *unc-71* (A Disintegrin And Metalloprotease protein) and *cam-1*/Ror kinase also exhibited defects in either primary neurite outgrowth or suppressed *kal-1gf)* ectopic branching. This is consistent with previous studies implicating both of these genes in shaping neuronal morphology (Huang et al. 2003; Kim et al. 2003; Kennerdell et al. 2009; Hayashi et al. 2009). Overall, these data indicate that multiple genes are required for both primary neurite outgrowth and *kal-1(gf)* ectopic axon branching in AIY neurons, in addition to EFN-4.

**TABLE 1.**
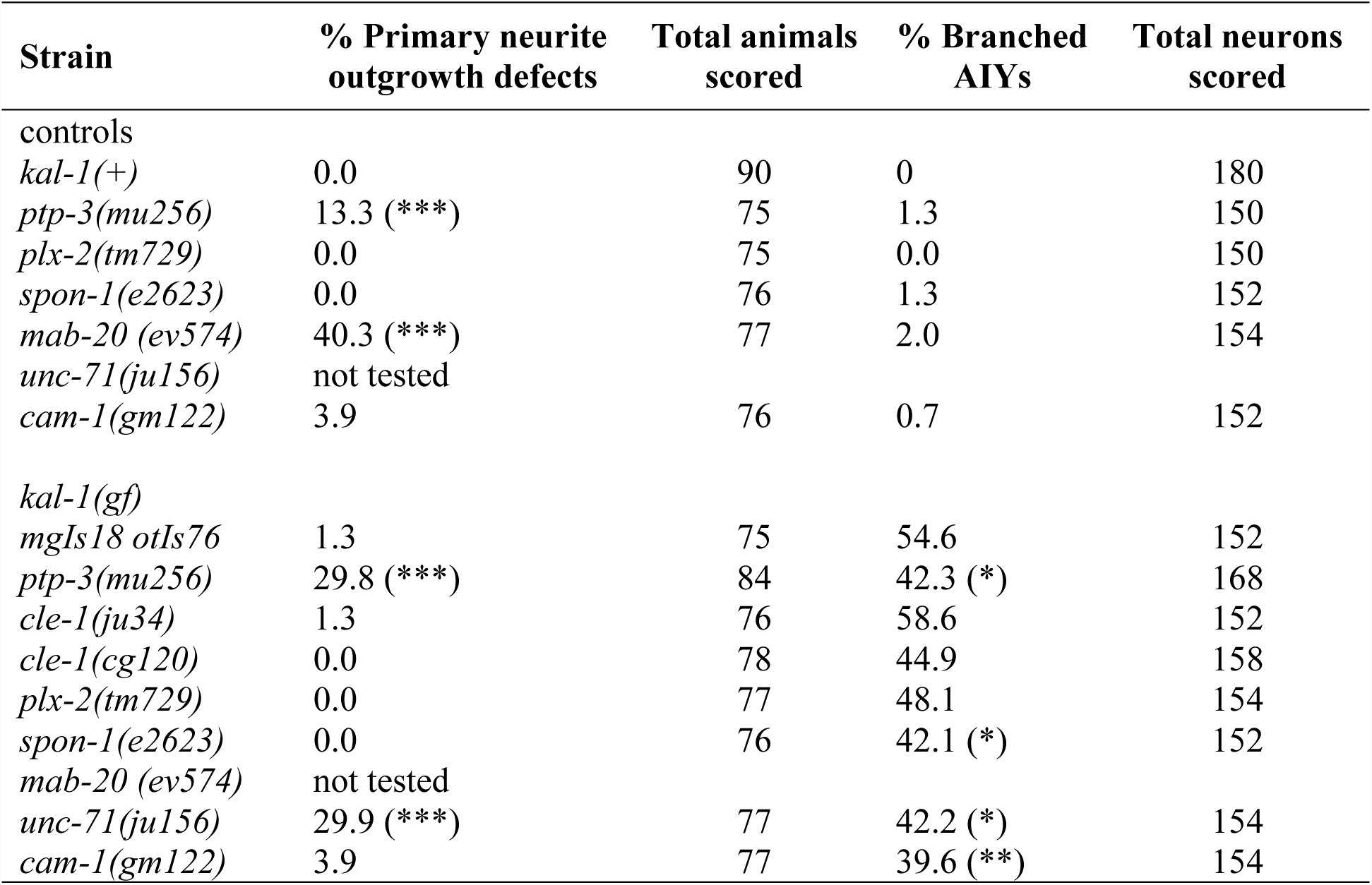
**Table 1 legend** Other neurodevelopmental genes examined during the course of this work. Control strains have *mgIs18 [P-ttx-3-GFP]* in the background. All *kal-1(gf)* strains contain *mgIs18 [P-ttx-3-GFP]otIs76[P-ttx-3-kal-1 P-unc-122-GFP].* Statistical significance was determined using the z-test. * p<0.05, ** p<0.01, *** p<0.001.

| Strain | % Primary neurite outgrowth defects | Total animals scored | % Branched AIYs | Total neurons scored |
| --- | --- | --- | --- | --- |
| controls |  |  |  |  |
| *kal-1(+)* | 0.0 | 90 | 0 | 180 |
| *ptp-3(mu256)* | 13.3 (***) | 75 | 1.3 | 150 |
| *plx-2(tm729)* | 0.0 | 75 | 0.0 | 150 |
| *spon-1(e2623)* | 0.0 | 76 | 1.3 | 152 |
| *mab-20 (ev574)* | 40.3 (***) | 77 | 2.0 | 154 |
| *unc-71(ju156)* | not tested |  |  |  |
| *cam-1(gm122)* | 3.9 | 76 | 0.7 | 152 |
| *kal-1(gf)* |  |  |  |  |
| *mgIs18 otIs76* | 1.3 | 75 | 54.6 | 152 |
| *ptp-3(mu256)* | 29.8 (***) | 84 | 42.3 (*) | 168 |
| *cle-1(ju34)* | 1.3 | 76 | 58.6 | 152 |
| *cle-1(cg120)* | 0.0 | 78 | 44.9 | 158 |
| *plx-2(tm729)* | 0.0 | 77 | 48.1 | 154 |
| *spon-1(e2623)* | 0.0 | 76 | 42.1 (*) | 152 |
| *mab-20 (ev574)* | not tested |  |  |  |
| *unc-71(ju156)* | 29.9 (***) | 77 | 42.2 (*) | 154 |
| *cam-1(gm122)* | 3.9 | 77 | 39.6 (**) | 154 |

### EFN-4 acts non-cell autonomously to promote AIY outgrowth and kal-1(gf) ectopic branching

As a GPI-linked cell surface ephrin, EFN-4 might act cell autonomously or non-autonomously from an adjacent cell to promote axon outgrowth. To address this, we used tissue-specific promoters driving expression of an *efn-4* cDNA to determine which tissue EFN-4 is required in for AIY primary neurite outgrowth versus *kal-1(gf)* dependent axon branching (Figure 4). Despite EFN-4’s extensive expression in the nervous system (Chin-Sang et al. 2002), both AIY-specific and pan-neural *efn-4* expression failed to rescue primary neurite outgrowth defects, indicating that EFN-4 may be functioning non-cell autonomously (Figure 4a).

**Figure 4.**
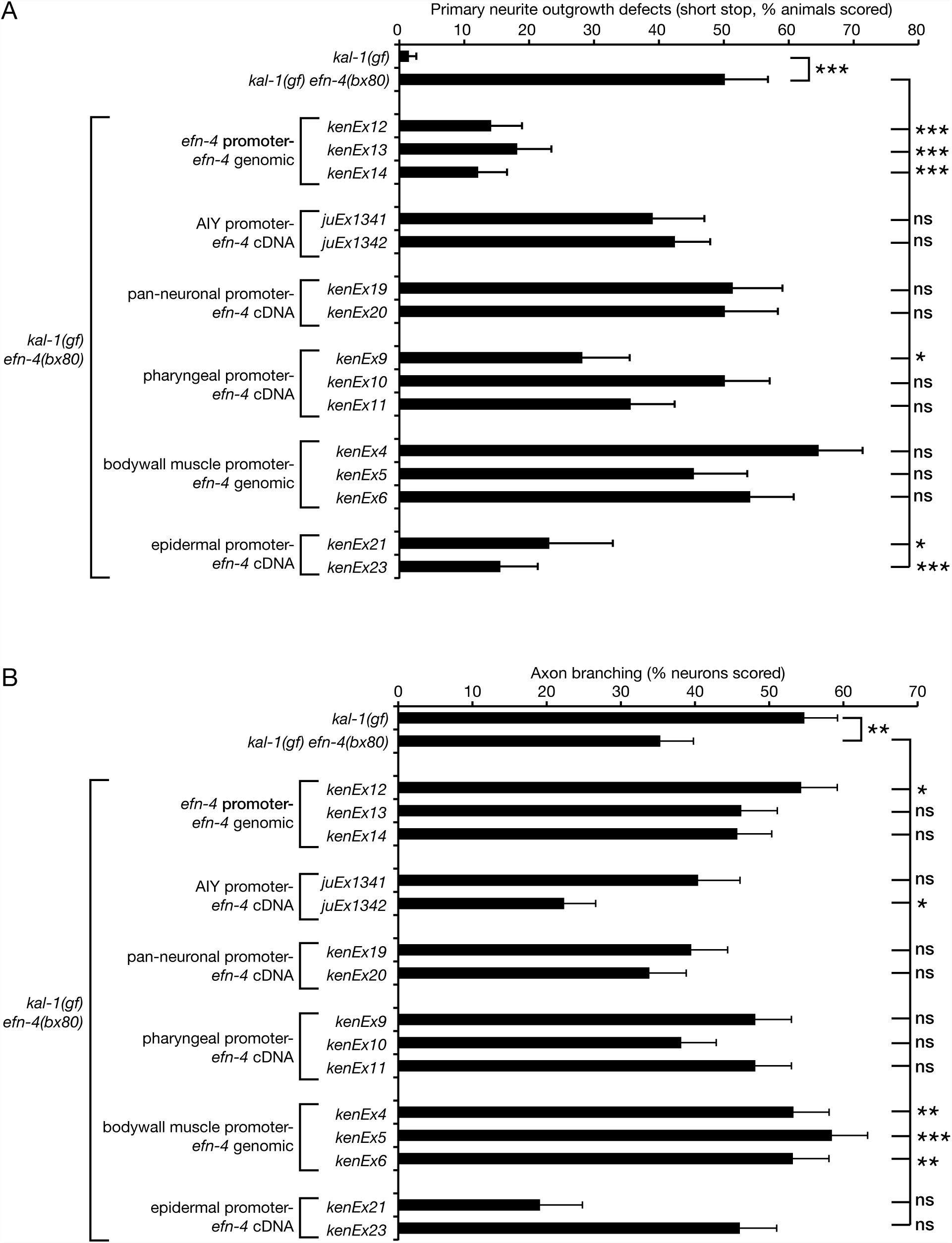
*efn-4* functions in the epidermis to promote AIY primary neurite outgrowth and in the body wall muscle to promote *kal-1(gf)-*induced ectopic axon branching. Tissue-specific *efn-4* transgenic lines were generated to establish what tissue *efn-4* is required in for (a) primary neurite outgrowth, and (b) ectopic neurite branching. *efn-4* genomic rescue data from Figures 1 and 3 are included for comparison. All strains contain *kal-1(gf)* and all rescue lines are in a *kal-1(gf) efn-4(bx80)* background. A minimum of two transgenic lines was assayed for every construct. Statistical significance was determined using the z-test. * p<0.05; ** p < 0.01; *** p < 0.001. Error bars are standard error of proportion.

The close juxtaposition of AIY cell bodies to pharyngeal, body wall muscle, and epidermal cells during neurite outgrowth suggested that one or more of these tissues could theoretically provide a non-cell autonomous cue for AIY neurite outgrowth. When we used the *myo-2* promoter to drive pharyngeal expression of an *efn-4* cDNA, only 1 out of 3 transgenic lines showed significant rescue of outgrowth defects (14/50 animals tested for that line). Expression of *efn-4* from a *myo-3* body wall muscle-specific promoter failed to show any rescue (0/3 lines tested, n=50 animals per line). However, expression of *efn-4* from an *elt-3* epidermal-specific promoter showed significant rescue (2/3 lines tested, n=21 and 49 animals per line respectively). Note that all three of the *P-elt-3-efn-4* lines showed high embryonic lethality, and one line strongly selected against epidermal *efn-4* expression. From this, we conclude that EFN-4 is required non-cell autonomously in the embryonic epidermis and possibly also the pharynx to promote AIY primary neurite outgrowth. We also conclude that embryonic development is highly sensitive to epidermal EFN-4 expression levels; the correct amount of EFN-4 in the appropriate tissue is sufficient to complete embryogenesis but over-expression can cause a neomorphic embryonic lethal phenotype.

We also examined whether tissue-specific expression of EFN-4 could rescue *kal-1(gf)* dependent axon branching (Figure 4b). AIY-specific expression of *efn-4* led to hyper-suppression of branching (1 out of 2 lines tested), where as pan-neural expression had no effect. Pharyngeal expression of *efn-4* showed a slight but not significant increase of branching in 2 out of 3 lines tested. Intriguingly, expression of *efn-4* in the body wall muscle fully rescued *kal-1(gf)* neurite branching in 3 out of 3 lines tested but did not rescue body morphology defects seen in *efn-4* mutants (data not shown). Finally, epidermal expression of *efn-4* was equivocal; one strain showed a decrease in branching whereas the other showed an increase, although neither were significantly different from the branching levels seen in *efn-4(bx80)* alone. From this, we tentatively conclude that EFN-4 expression from body wall muscle during early embryogenesis is required, in part, to promote *kal-1(gf)* axon branching. In addition, this suggests that *kal-1(gf)* ectopic branching is genetically separable from AIY primary neurite outgrowth.

### The Eph receptor VAB-1 plays a minor role in promoting AIY primary neurite outgrowth

If EFN-4 is required non-cell autonomously in the epidermis to promote AIY primary neurite outgrowth, then presumably it must function via one or more receptors on the AIY neurons. Previous work has identified multiple roles for the *C. elegans* Eph receptor VAB-1 during development, including embryonic morphogenesis, oocyte maturation, and amphid neuron axon guidance (George et al. 1998, Miller et al. 2003, Mohamed and Chin-Sang, 2006; Ikegami et al. 2012, Grossman et al. 2013). VAB-1 can bind EFN-4 in vitro making it an obvious candidate receptor (Wang et al 1999). We confirmed this finding using biolayer interferometry (BLI), a form of optical biosensing similar to surface plasmon resonance (Abdiche et al. 2008; Wilson et al. 2010). EFN-1::Fc, EFN-4::Fc or Fc control ligands were immobilized onto anti-human Fc capture probes then exposed to tissue culture supernatants containing the VAB-1 extracellular domain fused to alkaline phosphatase (VAB-1::AP). We found similar association curves for VAB-1::AP binding to both EFN-1 and EFN-4 (figure 5a) whereas Fc shows negligible association (the Fc binding data was used to normalize for negative drift hence data not shown). The dissociation curves for both EFN-1 – VAB-1 and EFN-4 – VAB-1 interactions were essentially flat, consistent with a very slow dissociation rate ( Figure 5b). These data reconfirm that EFN-4 binds VAB-1 in vitro.

**Figure 5.**
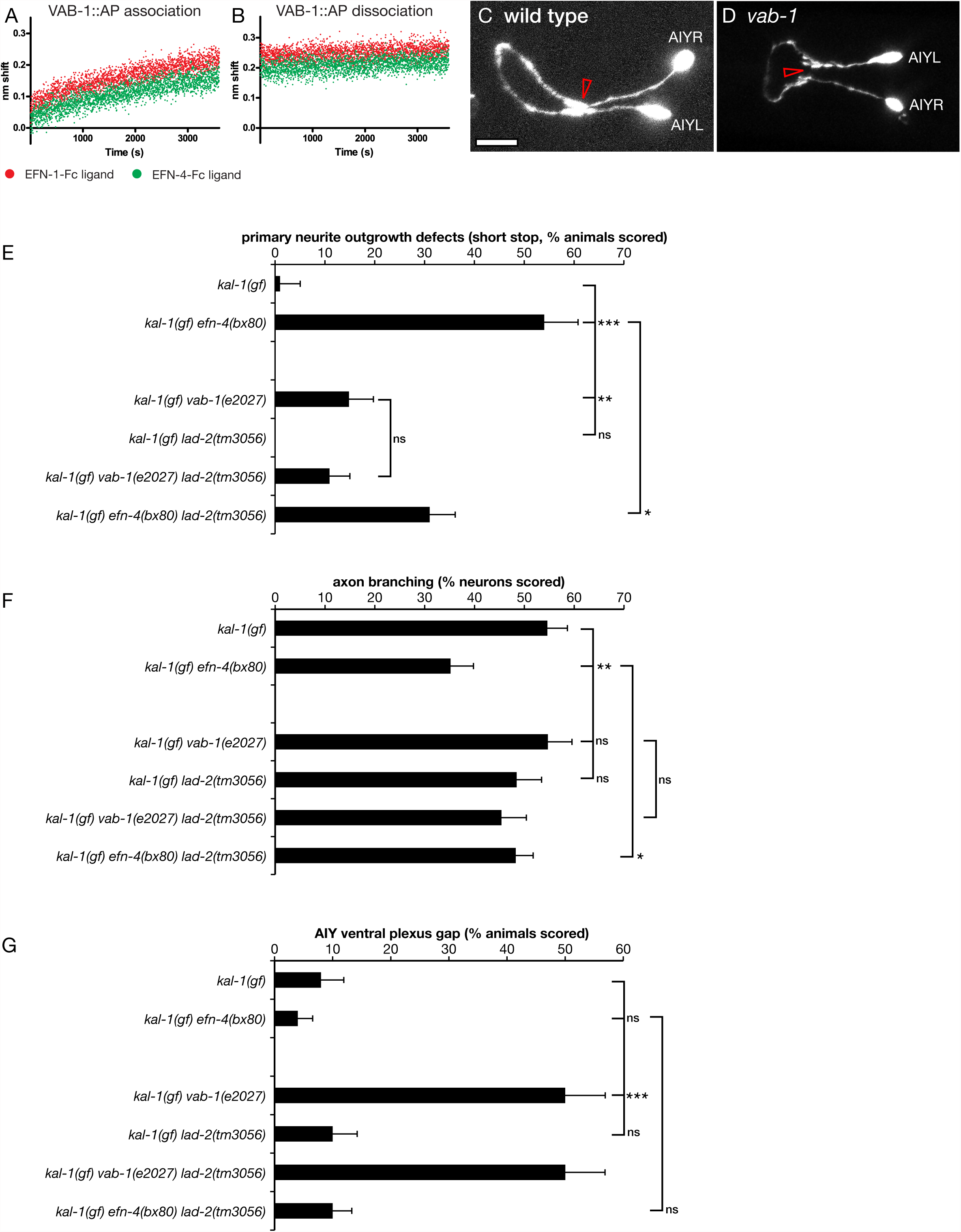
The Eph receptor *vab-1* promotes primary neurite outgrowth and ventral plexus contact in AIY neurons. Biolayer interferometry was used to demonstrate binding between the extracellular domain of VAB-1 fused to alkaline phosphatase and ephrins EFN-1 and EFN-4. (a) VAB-1::AP binding association phase. (b) VAB-1::AP dissociation phase. Binding curves are normalized against an Fc control to correct for instrument drift. (c, d) Confocal micrographs of AIY neurons in (c) wild type and (d) *vab-1(e2027)* mutants. Open arrow in (c) shows the ventral contact point in wild type AIY pairs. (d) *vab-1* mutants show a highly penetrant ventral gap where the AIYL/R cells fail to contact each other. Mutations in the candidate EFN-4 receptors *vab-1*/EphR and *lad-2*/L1CAM were assayed for (e) primary neurite outgrowth, (f) ectopic axon branching, and (g) ventral plexus contact. Statistical significance was determined using the z-test. * p<0.05; ** p < 0.01; *** p < 0.001. Error bars are standard error of proportion.

To ask whether VAB-1 plays any role in shaping AIY neuron development, we examined AIY morphology in a *vab-1(e2027)* null mutant background (George et al. 1998). In *vab-1(e2027)* animals, 15% of AIY neuron pairs (8/45) show short-stop defects during primary neurite outgrowth compared to 1% (1/76) of wild type worms (Figure 5e). However, this phenotype is significantly less than that seen in *efn-4(bx80)* animals (50%, 27/54). This suggests that VAB-1 plays a minor part in promoting AIY primary neurite outgrowth, and that it may function redundantly with one or more other receptors in this role. We were unable to assay whether *vab-1* and *efn-4* function together in a linear genetic pathway as *vab-1; efn-4* double mutant combinations show strongly synergistic embryonic lethality, thus preventing genetic analysis of AIY morphology in adult animals (Chin-Sang et al. 2002).

We also examined if VAB-1 plays a role in promoting the growth of *kal-1(gf)*-dependent ectopic neurites (Figure 5f). 55% of *kal-1(gf) vab-1(e2027)* neurons examined (58/106) showed ectopic branching, which is the same as that seen in *kal-1(gf)*. This suggests that VAB-1 plays no part in ectopic neurite branching and is consistent with EFN-4 having VAB-1-independent roles. This result also suggests there is an unidentified receptor for EFN-4 and provides further evidence that primary versus ectopic (secondary) neurite growth are genetically distinct processes.

### The Eph receptor VAB-1 is required for ventral plexus contact between AIY neurons

AIY neuron pairs have two contact points between the AIYL and AIYR cells; they initially touch each other in the ventral plexus then grow apart, wrapping around the left and right sides of the pharynx respectively, before contacting again on the dorsal side of the nerve ring via a gap junction (White et al. 1986). Although VAB-1 plays only a minor role in promoting primary neurite outgrowth, we find it plays a major role in organizing the plexus contact (Figure 5d, g). 50% of *vab-1(e2027)* animals (26/52) have a distinct gap between the AIY left and right cells at the plexus compared to 8% of wild type controls (4/49) and 4% of *efn-4* mutants (2/54). The functional importance of an AIYL/R plexus contact in the normal physiology of these cells is not known.

We also found that AIY neuron cell bodies were frequently displaced anteriorly in *vab-1* mutants compared to wild type, whereas *efn-4* cell bodies were often displaced posteriorly (data not shown). Despite this, we did not see any correlations between the location of the cell body and whether there were defects in primary neurite outgrowth or AIYL/R plexus contact. In addition, *vab-1* mutants frequently display variably penetrant head morphology defects, which could in principle influence the shape of neurons in the head region (George et al. 1998; Tucker and Han, 2008). However, we saw no correlation between head morphology defects and the presence of a ventral plexus gap between the AIYL and R neurons (5/9 animals with wild type head morphology had plexus gaps compared to 20/47 animals that displayed a morphological defect).

### The L1CAM LAD-2 may inhibit AIY primary neurite outgrowth

*efn-4* shows strong genetic interactions with the secreted semaphorin *mab-20*, and has been shown to bind in a complex with the semaphorin receptor PLX-2 (Nakao et al. 2007). MAB-20 and PLX-2 also exhibit genetic and physical interactions with the L1CAM LAD-2 (Wang et al. 2008). Our data shows that *mab-20* mutants exhibit strong AIY primary neurite shortstop defects, although PLX-2 has no obvious role in shaping AIY morphology (Table 1), raising the possibility that LAD-2 may be a receptor for EFN-4. We find that *lad-2(tm3056)* null mutants have no obvious defects in AIY primary neurite outgrowth when examined alone, but significantly suppress *efn-4* primary neurite defects in *efn-4; lad-2* double mutants (Figure 5e). Also, *vab-1; lad-2* double mutants are not significantly different from *vab-1* mutants alone. With regard to AIYL/R plexus contact, *lad-2* mutants are not significantly different from wild type and *vab-1; lad-2* double mutants display the same phenotype as *vab-1* mutants alone (27/54 *vab-1; lad-2* animals show a plexus gap phenotype compared to 26/52 *vab-1* mutants). These data suggest that LAD-2 may play a role in inhibiting AIY neuron primary outgrowth and that it functions antagonistically with EFN-4 in this process. We see no evidence of genetic interactions between *lad-2* and *kal-1(gf)* during ectopic neurite branching (Figure 5f). In addition, *lad-2* mutants have obvious AIYL/R plexus contact defects (Figure 5g). As there is no evidence for LAD-2 expression in AIY neurons (Wang et al. 2008), we consider it unlikely that LAD-2 is a cell autonomous receptor for EFN-4 during AIY neuronal development.

### EFN-4 may function in a parallel genetic pathway with 6-O-sulfated SDN-1/syndecan in AIY primary neurite outgrowth

Heparan sulfate proteoglycans have multiple roles in nervous system development and EphR/ephrin signaling has been shown to function in concert with HSPGs in mouse cortical neuron outgrowth (Van Vactor 2006; Irie et al. 2008). In *C. elegans*, the single-pass trans-membrane HSPG SDN-1/syndecan is required for neuroblast migration, axon outgrowth, axon guidance, and post-embryonic cell migration in multiple *C. elegans* CNS cell types (Rhiner et al. 2005; Hudson et al. 2006; Diaz-Balzac et al. 2014, Kinnunen 2014). In addition, LON-2/glypican has roles in the left/right guidance choices of DB motor neurons (Bulow et al. 2008). GPN-1/glypican is redundantly required for embryonic neuroblast migration but has no other reported function in *C. elegans* development (Hudson et al. 2006). We found that single mutants of the HSPGs *sdn-1*/syndecan, *gpn-1*/glypican and *lon-2*/glypican, had no significant effect on AIY primary neurite outgrowth (Figure 6). However, *sdn-1* loss-of-function mutations in combination with either *lon-2/*glypican or *gpn-1/*glypican showed low yet significant axon outgrowth defects, revealing redundant roles in AIY outgrowth among the HSPGs.

**Figure 6.**
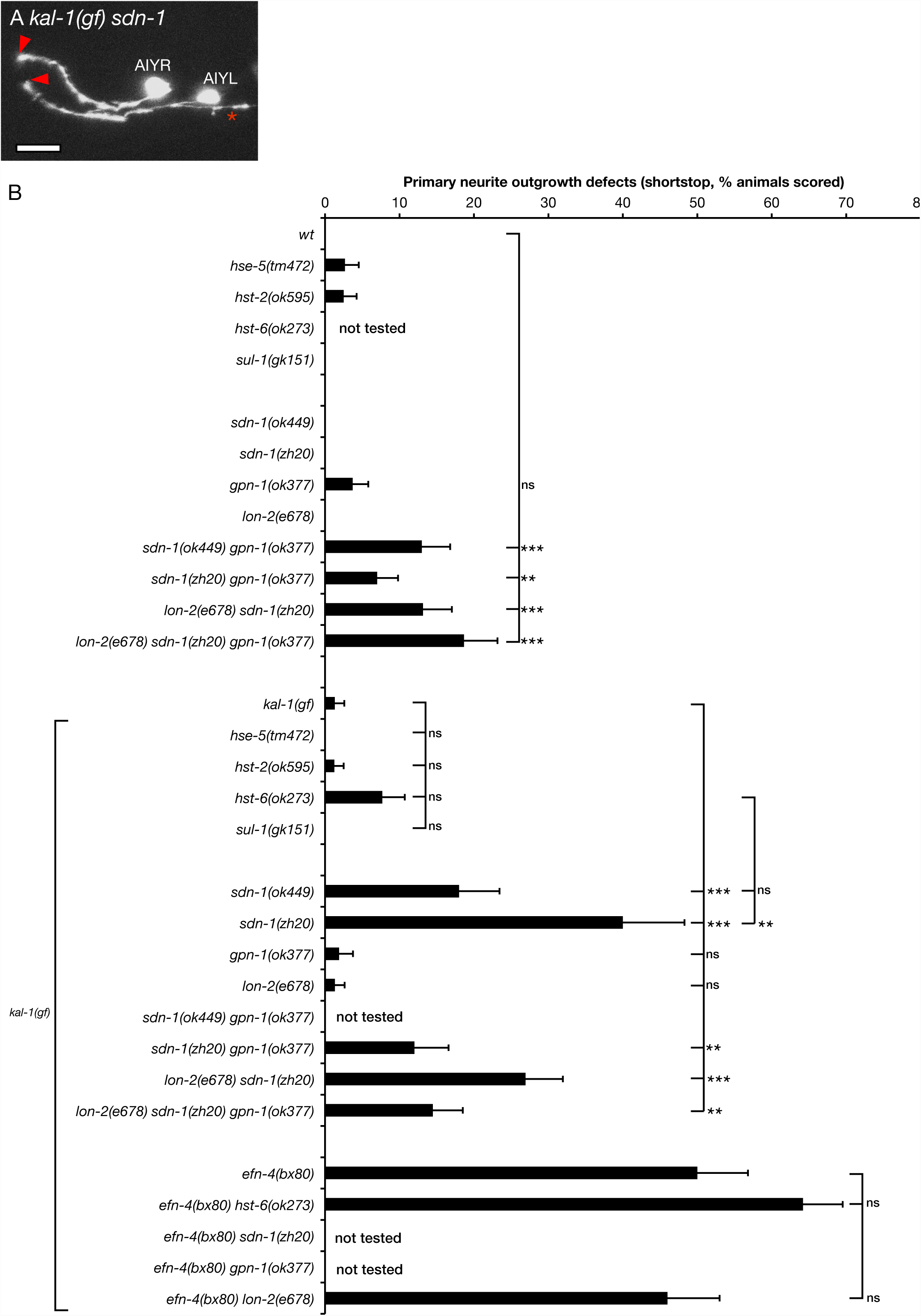
Heparan sulfate and heparan sulfate proteoglycans are required for AIY primary neurite outgrowth. (a) Confocal micrograph of AIY neuron morphology in a *kal-1(gf) sdn-1(zh20)* mutant. Arrowheads show premature termination of primary neurite outgrowth. Asterisk in shows a *kal-1(gf)* induced ectopic branch. Scale bar = 10 μm. (b) AIY primary neurite defects in HS-biosynthesis and HSPG core protein mutants assayed in wild type and *kal-1(gf)* mutant backgrounds. Statistical significance was determined using the z-test. * p<0.05; ** p < 0.01; *** p < 0.001. Error bars are standard error of proportion.

The *kal-1(gf)* background strongly sensitized AIY neurons to loss of *sdn-1*/syndecan. Primary neurite short-stop phenotypes were significantly enhanced to 18% for the *sdn-1(ok449)* weak loss-of-function allele and 44% for the *sdn-1(zh20)* null (Figure 6a, b). *sdn-1(ok449)* is an in-frame deletion of the *sdn-1* locus that removes the two known HS attachment sites yet retains the trans-membrane and C-terminus, including the conserved PDZ-binding domain (Rhiner et al. 2005; Dejima et al. 2014; Kinnunen 2014). The difference in phenotype penetrance observed in *sdn-1(ok449)* versus *sdn-1(zh20)* suggests that SDN-1 has both HS-dependent and independent roles in AIY primary outgrowth.

We also found that *kal-1(gf); sdn-1(zh20)* primary neurite defects were partially suppressed by both *lon-2* and *gpn-1* mutations. In addition, AIY-specific expression of *gfp*::*sdn-1* suppressed the *sdn-1* primary neurite shortstop phenotype (Figure 7, P < 0.05, 2 out of 3 lines tested), suggesting that SDN-1 functions cell autonomously. Conversely, expression of *gfp::gpn-1*from a *gpn-1* promoter was sufficient to increase primary neurite defects (1 out of 3 lines tested, P < 0.01). AIY-specific expression of *lon-2* showed a striking and significant increase in primary neurite shortstop defects (3 out of 3 lines tested, P < 0.001). Curiously, our *P-AIY-lon-2* transgenic lines showed highly penetrant loss of AIY cell bodies, which appeared to be transgene-specific (15% of *kenEx16* and 11% of *kenEx17* animals showed missing AIY cell bodies compared with 67% of *kenEx18* animals). In addition, we observed exuberant axon branching (multiple branches from a single neuron) and hyper-long axon branches (data not shown). Taken together, these data suggest that the shortstop phenotypes seen in *P-AIY-lon-2* animals may be influenced by the transgenic arrays titering out TTX-3, leading to neomorphic phenotypes similar to those seen in *ttx-3* weak loss-of-function mutations (Altun-Gultekin et al. 2001). Despite this, our data supports cell-autonomous yet antagonistic roles for SDN-1/syndecan and GPN-1/glypican in AIY primary neurite outgrowth, with possible contributions from LON-2.

**Figure 7.**
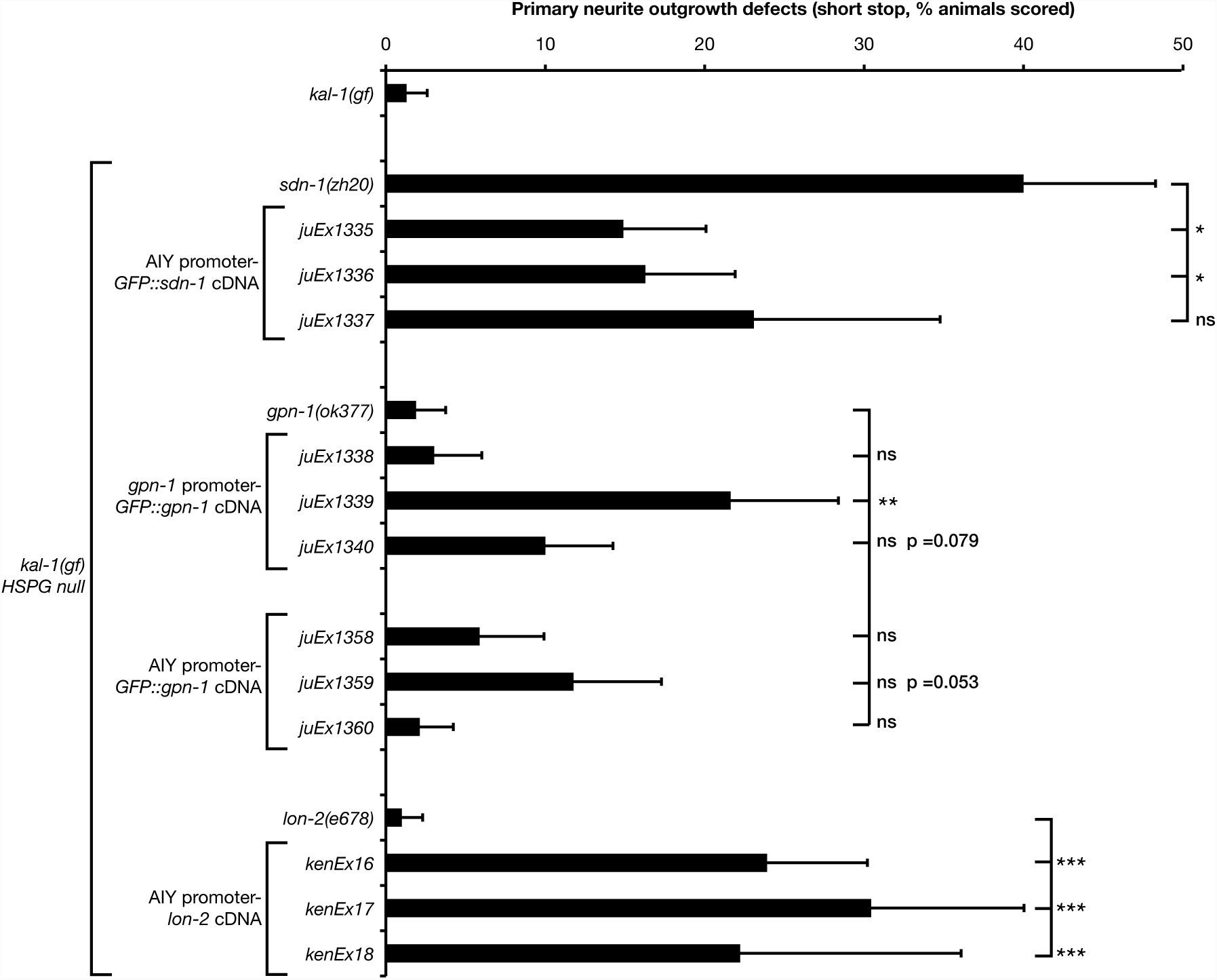
Cell autonomous expression of heparan sulfate proteoglycans reveals opposing roles for SDN-1 and glypicans in AIY primary neurite outgrowth. Primary neurite defects in *kal-1(gf) HSPG* null mutants bearing transgenic rescue arrays. *sdn-1(zh20)* animals contain *P-AIY-sdn-1::GFP* transgenes. *gpn-1(ok377)* animals contain *P-gpn-1*-*gpn-1::GFP* or *P-AIY-gpn-1::GFP* transgenes. *lon-2(e678)* animals contain *P-AIY-lon-2* transgenes. *sdn-1::GFP* and *gpn-1::GFP* transgenes were previously shown to be functional (Hudson et al. 2006). *HSPG* null mutant data from Figure 6 are included for comparison. Statistical significance was determined using the z-test. * p<0.05; ** p < 0.01; *** p < 0.001. Error bars are standard error of proportion.

Specific HS modifications have been shown to play distinct roles in *C. elegans* nervous system development (Bulow and Hobert 2004; Rhiner et al. 2005; Diaz-Balzac et al. 2014; Kinnunen 2014). We asked whether individual genes required for HS modification had any effect on AIY primary neurite outgrowth. Single mutants of *hse-5/*C5-epimerase, *hst-2*(2-O-sulfotransferase), *hst-6* (6-O-sulfotransferase) and *sul-1* (arylsulfatase) had either no effect on AIY primary neurites or were not significantly different when compared to wild type controls (Figure 6b). This was also the case when these mutants were examined in the *kal-1(gf)* background. Considering the phenotype seen in *sdn-1(ok449)* mutants, which is essentially an HS-minus version of the *sdn-1* gene, these data suggest that there is likely to be considerable redundancy between different HS sulfation patterns with regard to AIY primary neurite outgrowth.

The similarity in phenotypes between *efn-4* and *sdn-1* mutants suggests they may function in a linear genetic pathway to govern AIY primary neurite growth. However, *efn-4; sdn-1* double mutants exhibit strong synthetic lethality, making it difficult to directly address this question in adult worms (Hudson et al. 2006). As *hst-6* mutants show the strongest primary neurite phenotype (Figure 6, 8% outgrowth defects, 6/78 worms assayed), we asked whether *efn-4* had any genetic interaction with *hst-6* mutants. 64% of *efn-4; hst-6* double mutants showed primary neurite shortstop defects (52/81 animals scored), which is additive but not significantly different from *efn-4* mutants alone. *efn-4; lon-2* double mutants show 46% axon outgrowth defects (23/50 animals scored), which is not significantly different from *efn-4* alone (55%), although *lon-2* and *gpn-1* mutants additively and significantly suppress the *sdn-1* primary neurite outgrowth phenotype. From this, we conclude that HSPGs likely function in a parallel genetic pathway with *efn-4* during AIY primary neurite extension although other interpretations of these data are possible.

### EFN-4 acts in parallel to the KAL-1/HSPG pathway in ectopic axon branching

KAL-1-dependent ectopic branching is strongly suppressed by mutations in the HS modification genes *hst-2, hst-3.2, hst-6* and *hse-5*, indicating that this phenotype is an HS dependent process (Bulow et al. 2002; Tecle et al. 2013). We confirmed some of these observations using deletion alleles of *hst-2, hst-6* and *hse-5* (Figure 8b). We find that null mutations of HS modifier genes suppress *kal-1(gf)* branching to about 15-20% of AIY neurons, which is similar to that reported by others (Bulow et al. 2002).

**Figure 8.**
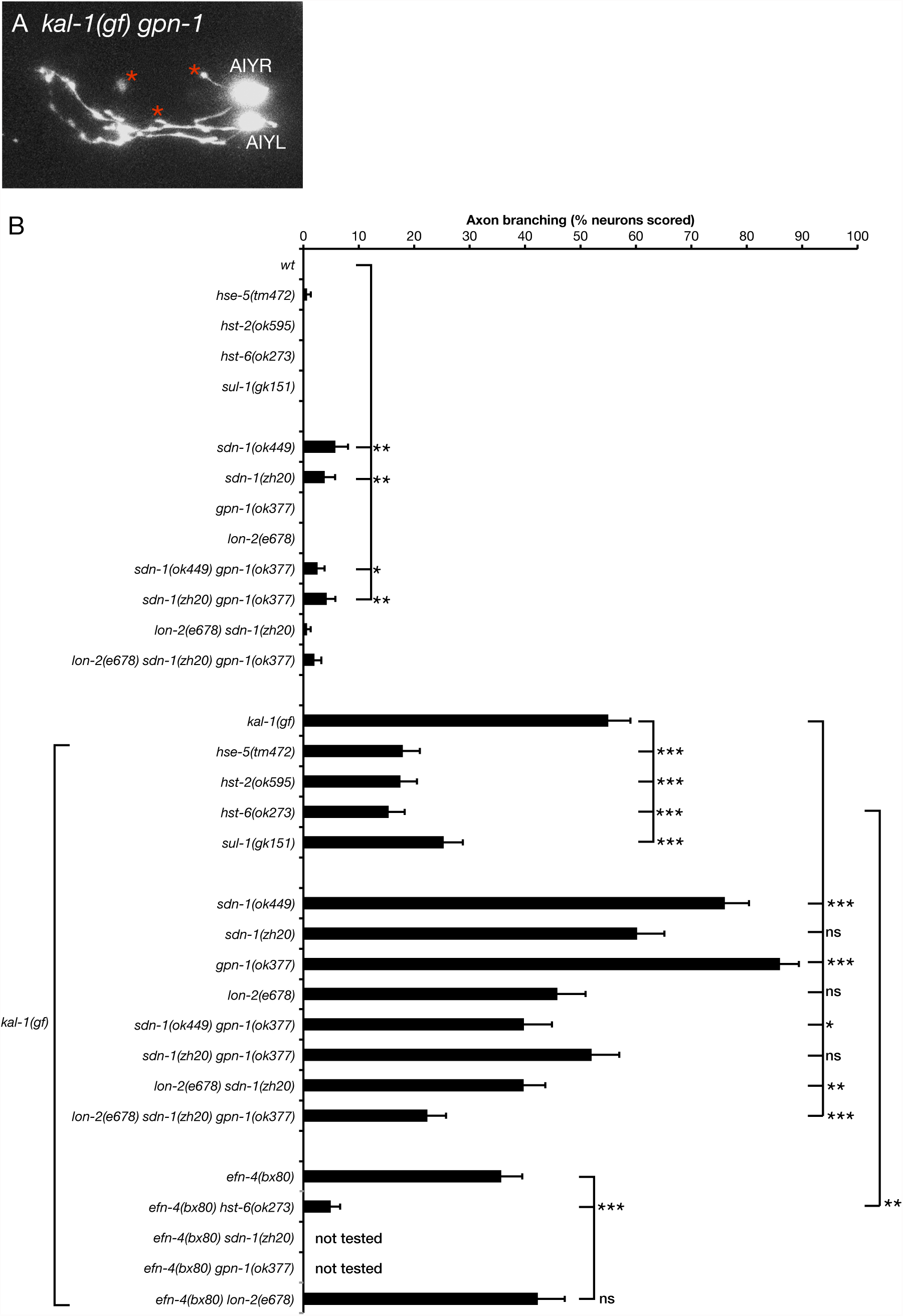
Heparan sulfate and heparan sulfate proteoglycans are required for *kal-1(gf)* induced ectopic AIY neuron branching and function in parallel with EFN-4. (a) Confocal micrograph of AIY neurons in a *kal-1(gf) gpn-1(ok377)* mutant. (b) AIY ectopic branching defects in HS-biosynthesis and HSPG core protein mutants assayed in wild type and *kal-1(gf)* mutant backgrounds. Pertinent double mutant combinations with *efn-4(bx80)* are also included. Statistical significance was determined using the z-test. * p<0.05; ** p < 0.01; *** p < 0.001. Error bars are standard error of proportion.

The partial suppression of *kal-1(gf)* ectopic neurite branching and our data indicating that *efn-4* may function in parallel with HSPGs during AIY primary neurite outgrowth suggests *efn-4* may also function in parallel with HSPGs during axon branching, similar to its role in embryogenesis (Hudson et al. 2006). Consistent with this interpretation, *efn-4; hst-6* double mutants caused significant suppression of *kal-1(gf)* branching to background levels (5%) compared to *hst-6* alone (15%, Figure 8b). We conclude that EFN-4 and HSPGs have parallel and partly redundant roles in KAL-1-induced ectopic neurite branching.

We also tested requirements for another HS modifier enzyme not previously examined for its roles in *kal-1(gf)* branching. *sul-1* encodes the *C. elegans* ortholog of mammalian extracellular sulfatase Sulf1 (Ai et al. 2003; Nawroth et al. 2007). The deletion mutation *sul-1(gk151)* is predicted to eliminate most of the conserved functional domains of SUL-1 and thus cause either complete or strong loss of function. *sul-1(gk151)* animals were viable and fertile. We found that *sul-1* strongly suppressed *kal-1(gf)* branching (25%). As SUL-1 is predicted to remove 6-O-sulfate groups from sulfated N-acetylglucosamine residues of HSPGs, this result suggests that KAL-1 may be unable to interact with over-sulfated HS.

We next examined the roles of HSPG core proteins in AIY neuron ectopic branch formation (Figure 8). Our previous analysis of embryonic morphogenesis identified the cell surface HSPGs syndecan SDN-1 and glypican GPN-1 as acting redundantly in a common pathway with KAL-1 (Hudson et al. 2006). In contrast to the requirement for SDN-1 in AIY neuron primary neurite outgrowth, we found that *sdn-1(ok449)* mutants showed increased ectopic branching, although this phenotype was not observed in *sdn-1(zh20)* null mutants. This was surprising because SDN-1 appears to be the most abundant HSPG in the *C. elegans* nervous system; *sdn-1(ok449)* mutants display dramatically reduced levels of neuronal anti-stub 3G10 immunoreactivity, an antibody that detects HS attachment sites on HSPG core proteins following heparatinase digestion (Minniti et al. 2004). Similarly, *gpn-1(ok377)* mutants also showed increased ectopic branching (Figure 8a, b), suggesting that GPN-1 plays an antagonistic role in this phenotype. We then tested whether SDN-1 and GPN-1 might act redundantly in KAL-1-induced branching. However, *sdn-1 gpn-1* double mutants either showed no suppression (*zh20 ok377*, 52%) or only partial suppression (*ok449 ok377*, 40%, P < 0.05) of *kal-1(gf)* branching. We therefore tested whether the second glypican, LON-2, might be a third redundant component of the KAL-1/HSPG network in axon branching. *lon-*2 single mutants displayed partial suppression of *kal-1(gf)* (45%, not significant), and this was not further enhanced in *lon-2 sdn-1,* indicating that LON-2 contributes to KAL-1-induced branching. *lon-2 sdn-1 gpn-1* triple null mutants were viable and showed stronger suppression of *kal-1(gf)* branching to levels similar to that of *kal-1(gf) hst-6(ok273)* (22% compared to 15%, P = 0.11). We conclude that all three *C. elegans* cell surface HSPGs have partly redundant roles in KAL-1-induced ectopic neurite branching.

We used genomic and tissue-specific rescue experiments to clarify the roles of SDN-1, GPN-1 and LON-2 in *kal-1(gf)* ectopic branching (Figure 9). AIY-specific expression of a *gfp::sdn-1* cDNA strongly suppressed ectopic branching phenotypes, suggesting that SDN-1 functions cell autonomously (3 out of 3 lines, P < 0.001). A *gpn-1::GFP* genomic clone also significantly suppressed the *gpn-1(ok377)* hyper-branching phenotype back to levels seen in *kal-1(gf)* animals alone (3 out of 3 lines). This result was mirrored by AIY-specific *gpn-1* transgenic arrays, suggesting that GPN-1 also functions cell autonomously. Finally, genomic expression of *lon-2* showed significant increases in ectopic branching (one out of three lines tested), as did AIY-specific expression of *lon-2* (2 out of 3 lines, P < 0.05), although this AIY-specific data should be viewed with caution based on the neomorphic missing cell phenotype mentioned previously. Epidermal expression of *lon-2* had no effect on ectopic branching but did rescue *lon-2* mutant body length defects (Gumienny et al. 2007). Taken together, these data point to cell autonomous, negative regulatory roles for both SDN-1 and GPN-1 in *kal-1(gf)* ectopic branching, with a possible role for LON-2 as a positive regulator of this phenotype. While this result appears paradoxical when compared to the robust suppression seen in *sdn-1 gpn-1 lon-2* triple mutants, the *kal-1(gf)* branching phenotype is driven by over-expression of KAL-1, which is a known HS-binding protein (Bulow et al. 2002; Hudson et al. 2006). We conclude that KAL-1-dependent axon branching is likely to be caused by an imbalance between KAL-1 and its negative regulators SDN-1 and GPN-1, leading to over-activation of the KAL-1 pathway and exuberant axon branch formation. LON-2 may function downstream or in parallel with KAL-1 in this process, based on the ability of genomic LON-2 to increase ectopic branching. Finally, the robust branching suppression found in HS modification mutants underlines the importance of distinct HS sulfation patterns in this process, suggesting that a particular HS sulfation motif is required for KAL-1 function.

**Figure 9.**
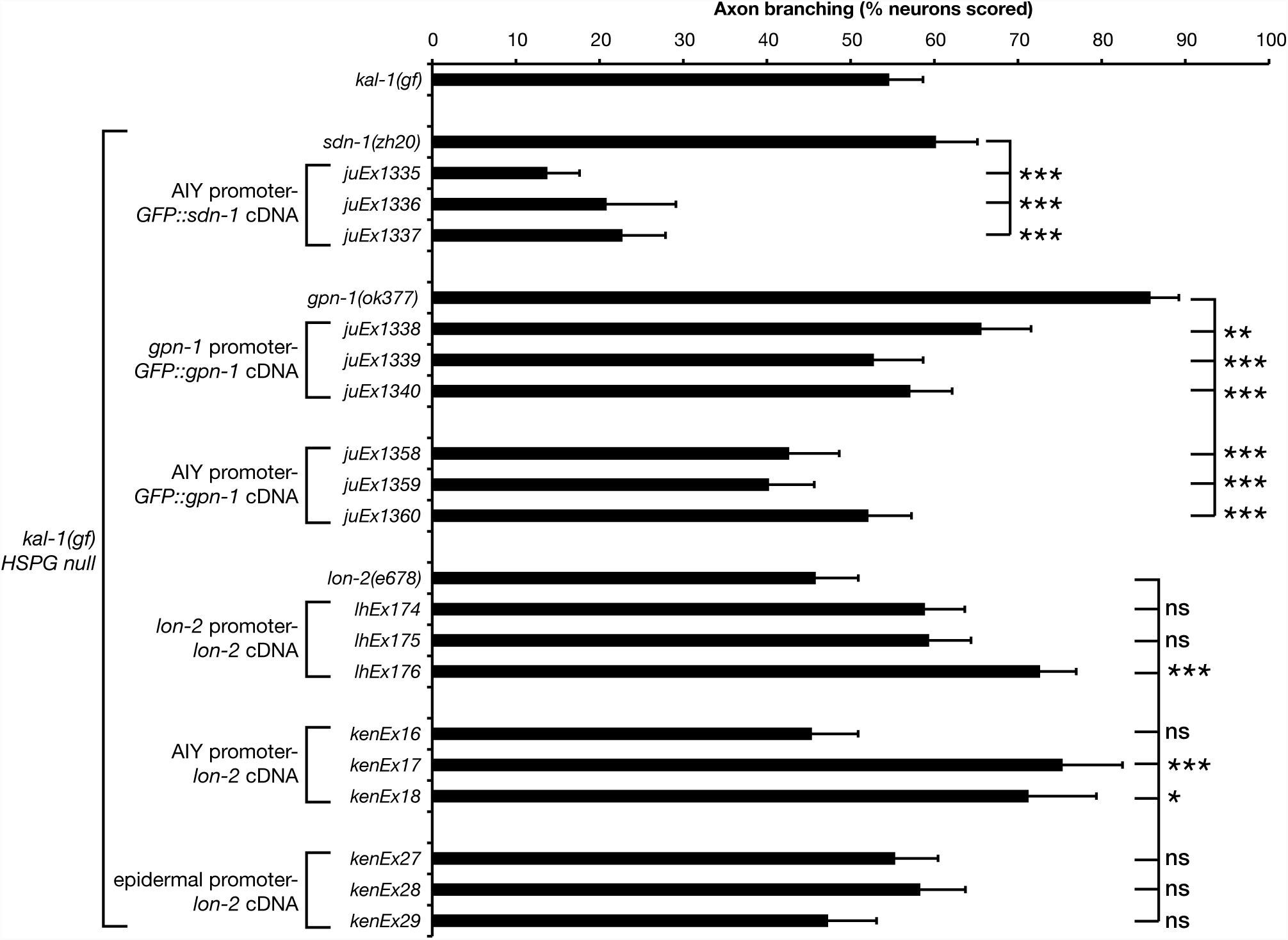
Over-expression of HSPG core proteins SDN-1 and GPN-1 can suppress *kal-1(gf)* induced ectopic AIY neuron branching. Ectopic branching in *kal-1(gf) HSPG* null mutants bearing transgenic rescue arrays. *kal-1(gf)* ectopic branching defects in *sdn-1(zh20)* animals containing *P-AIY-sdn-1::GFP* transgenes; *gpn-1(ok377)* animals containing genomic-*gpn-1::GFP* and *P-AIY*-*gpn-1::GFP* transgenes; and *lon-2(e678)* animals containing genomic-*lon-2*, *P-AIY*-*lon-2* and epidermis-specific-*lon-2* transgenes. *sdn-1(zh20), gpn-1(ok377)* and *lon-2(e678)* data from Figure 8 are included for comparison. Statistical significance was determined using the z-test. * p<0.05; ** p < 0.01; *** p < 0.001. Error bars are standard error of proportion.

## DISCUSSION

The data presented here identify a novel role for the *C. elegans* ephrin EFN-4 in shaping neuronal morphology. EphR – ephrin signaling functions primarily in a contact-dependent manner during nervous system development. For instance, counter-gradients of EphRs and ephrins presented in the mammalian superior colliculus provide axon targeting information for retinal ganglion cell growth cones navigating from the developing retina (Flanagan 2006). In this context, the target neuronal tissue presents positional cues to a navigating neural cell type. Also, EphRs and ephrins expressed in developing rhombomeres help drive their segregation into individual units during vertebrate hindbrain development (Mellitzer et al. 1999). Again, this EphR - ephrin interaction is solely within the nervous system. Our data indicate a novel role for EFN-4 where it acts non-cell autonomously from the epidermis to provide an axon outgrowth cue to AIY neurons.

### EFN-4 as an axon-targeting cue

While the non-cell autonomous role of EFN-4 in AIY axon outgrowth is novel, there is extensive evidence for non-cell autonomous roles of other molecules in nervous system development. For instance, UNC-6/netrin expression in the ventral epidermis provides pioneer growth cues to motorneurons during development of the nervous system (Wadsworth et al. 1996). In addition, SLT-1 expression from dorsal body wall muscles guides AVM neuron axon path finding (Hao et al. 2001). While the netrin, slit/robo and EphR/ephrin pathways have key roles in many aspects of nervous system development, netrin and slit are both secreted guidance cues, whereas ephrins are anchored to the cell membrane. In addition, AIY axon trajectories in *efn-4* mutants are normal in that both AIYL/R axons enter the nerve ring at the correct location, but stop short of their dorsal target. This suggests that EFN-4 is required, in part, to promote the final step of AIY primary neurite extension. Our previous work demonstrated a distinct region of EFN-4 expression in the anterior epidermis of lima-bean stage embryos (Nakao et al. 2007). We speculate that this abrupt EFN-4 expression boundary may provide the guidance information required for AIY primary neurites to complete their dorsal migration.

Our rescue data also suggest that EFN-4 presented by the nascent pharynx may be partially required for AIY axon outgrowth (Figure 4). In support of this, EFN-4::GFP expression is observed in pharyngeal tissue (Chin-Sang et al. 2002). Also, growth cones have already begun to migrate and encircle the developing pharynx at lima-bean stage, which is the earliest onset of *P-ttx-3-GFP* expression in AIY interneurons (Christensen et al. 2014 plus our unpublished observations). While our most robust rescue data is driven from an *elt-3* epidermal promoter, it may be that EFN-4 presented from the pharynx can functionally compensate for epidermal EFN-4, bearing in mind that both tissues are close enough for cell-cell contact during this stage of AIY neuron development.

### Canonical and non-canonical receptors for EFN-4

If EFN-4 is providing non-cell autonomous growth cues to AIY axons, then there must be at least one receptor for EFN-4 being expressed on the AIY interneurons. The *C. elegans* genome contains only one predicted canonical EphR gene, *vab-1*. We find that *vab-1* mutations only partially phenocopy *efn-4* defects in axon outgrowth, and instead, exhibit a novel, highly penetrant ventral contact defect between the AIYL and AIYR cells. VAB-1 has a previously defined role in ventral midline axon fascicle maintenance, where it functions with EFN-1 and the cell surface immunoglobulin superfamily protein WRK-1 (Boulin et al. 2006). In *C. elegans* embryos, EFN-1 and WRK-1 expression in the embryonic motor neuron cell bodies function as a ventral midline pioneer cue for PVQ, HSN, and command interneurons. Expression of VAB-1 in the HSN neurons alone is sufficient to rescue *vab-1* HSN guidance defects. Also, VAB-1 can physically interact in a complex with WRK-1 and EFN-1. Whether WRK-1 or EFN-1 has roles in AIYL/R plexus contact remains to be determined.

Previous genetic data suggest that EFN-4 may represent a novel, non-canonical member of the *C. elegans* ephrin family; *vab-1* mutants have striking head and tail morphology defects, a phenotype shared with *efn-1* mutants, whereas *efn-4* mutants display only partially penetrant mid-body and tail defects (George et al. 1998, Chin-Sang et al. 1999; Chin-Sang et al. 2002). In addition, *vab-1; efn-4* double null mutants exhibit synthetic embryonic lethality, suggesting that EFN-4 and VAB-1 each have one or more novel interaction partners during *C. elegans* embryogenesis. In addition to VAB-1’s previously demonstrated binding interaction with WRK-1, it has also been shown to physically bind to SAX-3/robo (Boulin et al. 2006; Ghenea et al. 2005). Also, *sax-3* mutants share similar head morphology and axon outgrowth defects with *vab-1* mutants (Hao et al. 2001; Ghenea et al. 2005). Curiously, the highly penetrant body morphology defects found in *sax-3* mutants is at odds with those presented by *slt-1* mutants, which are essentially wild type in appearance. Taken together, these observations suggest caution when drawing conclusions based on genetic interactions alone.

Our optical biosensing data have helped to clarify the conundrum presented by *vab-1* and *efn-4*’s dissimilar mutant phenotypes. We find that the VAB-1 extra-cellular domain can bind rapidly to both EFN-1 and EFN-4 in vitro. This interaction is qualitatively high affinity in that we see no dissociation of VAB-1 from the EFN-1 and EFN-4 coated biosensors (Figure 5a, b). Our data support previous findings, demonstrating that Fc-fusion proteins of EFN-1, 2, 3, and 4 can immunoprecipitate VAB-1 expressed in transiently transfected COS-1 cells (Wang et al. 1999). In addition, the intimate juxtaposition of EFN-4-expressing epidermal cells with VAB-1 expressing ventral neurons suggests that a physical interaction can occur between EFN-4 and VAB-1 during embryogenesis (George et al. 1998; Chin-Sang et al. 1999; Chin-Sang et al. 2002; Nakao et al. 2007). While our data suggest a role for VAB-1 in promoting EFN-4-dependent primary neurite extension, we also find that at least one other receptor is required for this process. Although the L1CAM LAD-2 may have a modest role in inhibiting EFN-4- dependent primary neurite outgrowth, it is unlikely to be the EFN-4 receptor required for neurite extension, raising the possibility that one or more novel EFN-4 receptors exist.

### EFN-4 has genetically distinct roles in kal-1 gain-of-function ectopic branching

In addition to defining a role for EFN-4 in AIY primary neurite outgrowth, we find that EFN-4 is also required, in part, for *kal-1(gf)*-dependent ectopic neurite branching (Bulow et al. 2002). While the correlation between primary versus secondary neurite growth might seem obvious at first, we find that these two processes are genetically distinct. Epidermal expression of EFN-4 does not rescue ectopic neurite outgrowth, whereas muscle expression of EFN-4 does. Cell tracking data and *myo-3-GFP* transgene studies reveal that myoblasts are born adjacent to the AIY interneurons, making it possible that muscle expression of EFN-4 is at least partially required for ectopic secondary neurite outgrowth (Bao et al. 2006; Tucker and Han 2008). In addition, UNC-52/perlecan, which is exclusively expressed in body wall muscle, has roles in ectopic neurite branching (Diaz-Balzac et al. 2014). The robust branching rescue seen in *P-myo-3-efn-4* lines is somewhat at odds with the weak restoration of phenotype seen in *efn-4* genomic rescues, but could be explained by the mosaic nature of *C. elegans* transgenic expression and the lack of an epidermal-specific marker in our genomic rescue arrays.

### SDN-1 and glypicans have opposing roles in both primary neurite outgrowth and ectopic branching

Previous studies have demonstrated the importance of HSPGs in nervous system development, where they have roles in TGFβ, Hedgehog, Wnt and Slit/robo signaling (Perrimon and Hacker 2004; Yan and Lin 2008; Schaefer and Schaefer 2009; Taneja-Bageshwar and Gumienny 2012). Despite this, little is known about how HSPGs and EphR/ephrin pathways function in concert. Studies in mouse reveal that both HSPGs and ephrin-A3 function together during mouse cortical axon outgrowth (Irie et al. 2008). Also, retinal ganglion cells require HSPGs and Slit for accurate navigation at the optic chiasm (Plump et al. 2002; Pratt et al. 2005). Hence retinal ganglion cells require both HS and EphR/ephrin cues at distinct phases of neural development. Our data suggest that EFN-4 and HSPGs are both required during AIY primary neurite outgrowth. The *kal-1(gf)* background employed in much of our work sensitizes the AIY neurons to *sdn-1* loss-of-function, revealing primary neurite outgrowth defects similar to those seen in *efn-4* mutations. Also, cell-autonomous expression of *sdn-1* in the AIY neurons strongly suppresses *sdn-1* primary neurite extension defects. This suggests that SDN-1/syndecan has a cell-autonomous role in promoting AIY primary neurite outgrowth, which is in broad agreement with previously published data on the role of SDN-1 in other neuron cell types (Minniti et al. 2004; Rhiner et al. 2005; Bulow et al. 2008; Kinnunen 2014).

Intriguingly, *sdn-1* phenotypes are suppressed by both *lon-2* and *gpn-1* mutations, whereas cell-autonomous expression of GPN-1 or LON-2 causes increased primary neurite outgrowth defects. *gfp::gpn-1* is expressed in both neural and non-neuronal cells (M. L. H., unpublished observations) whereas LON-2 functions from the epidermis to negatively regulate TGFβ signaling (Gumienny et al. 2007). Our tissue specific expression data strongly suggest a cell-autonomous role for GPN-1 glypican in negatively regulating KAL-1(gf) neurite extension. Also, the strong increase in *P-AIY-lon-2* dependent primary neurite defects suggests that LON-2 may also a negative regulator of outgrowth. However, we caution against a simple interpretation of this. First, there is no evidence of LON-2 expression in the nervous system (Gumienny et al. 2007). Second, we suspect that the *P-AIY*-*lon-2* lines may be exhibiting *ttx-3-*like weak loss-of-function phenotypes that are masking the underlying data (Altun-Gutelkin et al. 2001). Despite these observations, our *lon-2* loss-of-function data is robust and continues to support a role for LON-2 during AIY primary neurite outgrowth and branching, consistent with previous work demonstrating part-redundant roles for LON-2 in nervous system development (Diaz-Balzac et al. 2013; Kinnunen 2014).

We also investigated the role of specific HS modifications in AIY primary neurite outgrowth and branching. We find that loss of individual HS modification enzymes leads to very weak phenotypes in primary neurite extension yet very strong phenotypes in ectopic neurite formation. This suggests that HS plays only a minor role in primary neurite outgrowth. However, the HS-less *sdn-1(ok449)* allele shows a stronger phenotype that individual HS-modification mutants, indicating that HS-side chains are required for this process. It may be that there is sufficient redundancy in HS sulfation and modification patterns that different HS motifs can functionally compensate for one another during AIY primary neurite extension, similar to that reported for the genetic redundancy between *hst-2*, *hst-6*, and *hse-5* during DA and DB motorneuron outgrowth (Bulow et al. 2008).

The ectopic branching phenotype seen in *kal-1(gf)* animals represents a Kallmann syndrome model, and was originally used as a genetic screening background to identify novel genes that may be involved in this neurodevelopmental disorder (Bulow et al. 2002; Diaz-Balzac et al. 2014). We confirmed and extended these findings, revealing that multiple enzymes in the HS biosynthesis pathway are required for ectopic branch formation, including the arylsulfatase ortholog SUL-1. The role of HSPG core proteins in *kal-1(gf)* ectopic branching broadly parallels their roles in primary neurite outgrowth, although there is considerable redundancy in their interplay. Our data strongly support cell-autonomous antagonistic roles between SDN-1 and GPN-1, functioning in *cis* in both primary neurite extension and ectopic branching. This is in contrast to the role of *dally*/glypican in *Drosophila,* which functions antagonistically and in *trans* with *Dsdn*/syndecan and LAR/receptor tyrosine phosphatase during neuromuscular junction development (Johnson et al. 2006).

We also uncovered an epistatic relationship between *lon-2* and *sdn-1* during ectopic branch formation, where *lon-2* mutants consistently suppress *sdn-1* dependent defects. Although this suppression is not significant in *lon-2 sdn-1* double mutants, it is significant in *lon-2 sdn-1 gpn-1* triple mutants, suggesting partial redundancy between *lon-2* and *gpn-1* in this process.

The branching suppression seen in our HSPG triple mutant is not significantly different from that seen in our strongest HS-modifier suppressor mutant, *hst-6(ok273)*, confirming previously data on the importance of 6-O-sulfated HS to *kal-1(gf)* ectopic branching (Bulow et al. 2002). As our *hst-6* phenotype is no different from the *sdn-1 gpn-1 lon-2* triple mutant data, we cautiously suggest that no other HSPGs are involved in this process. This is in contrast to data presented by others which indicate a role for the extracellular matrix HSPG UNC-52/perlecan in *kal-1(gf)* branching (Diaz-Balzac 2014). We did not examine *unc-52* mutations because null alleles of this gene are embryonic lethal (Rogalski et al. 1993; Moerman et al. 1996). As such, we cannot discount a part-redundant role for UNC-52 in promoting *kal-1(gf)* branching. Overall, our data indicate that SDN-1, GPN-1 and LON-2 function together to govern *kal-1(gf)* ectopic branching, and that this process is intrinsically HS-dependent.

### EFN-4 and HSPGs function in parallel pathways to promote *kal-1(gf)* ectopic branching

Our previous analysis of KAL-1, SDN-1/syndecan and GPN-1/glypican in embryogenesis indicated that all three genes function in parallel with EFN-4 (Hudson et al. 2006). In this study, we took advantage of the *kal-1(gf)* ectopic branching phenotype to further explore the genetic relationship between these genes. Unfortunately, the strong synthetic lethality seen between *sdn-1* and *efn-4* mutants makes it difficult to directly address their interplay during AIY neuron primary neurite outgrowth and ectopic branching. However, we have indirect evidence that they function in parallel pathways. In ectopic neurite branching, EFN-4 is clearly functioning in parallel with a pathway requiring 6-O-sulfated HS, as *efn-4; hst-6* double mutants show the strongest axon branching suppression. Second, *lon-2* shows no genetic interaction with *efn-4*, beyond a modest increase in embryonic lethality (M.L.H., unpublished observations), whereas *lon-2* can suppress *sdn-1* phenotypes. Taken together, we conclude that EFN-4 and HSPGs function in parallel to promote *kal-1(gf)* ectopic neurite branching (and probably AIY primary neurite outgrowth also), although other interpretations of these data are possible.

#### A possible role for multiple ephrins in shaping a single neuron?

Overall, our data point to a model where EFN-4 functions in the epidermis to promote AIY primary neurite outgrowth, possibly by binding to VAB-1 and one or more additional co-receptors (Figure 10). This interaction is in parallel with HSPGs, which function cell autonomously in primary neurite outgrowth. In addition, VAB-1 has a non-EFN-4 dependent role in shaping the ventral contact point between the AIYL and AIYR cells. Whether this is a cell autonomous or non-cell autonomous role for VAB-1 remains to be discovered. Intriguingly, an EFN-1::GFP reporter gene is expressed in the AIY interneurons, suggesting that VAB-1 may function non-cell autonomously to drive AIYL/R contact (Grossman et al. 2013). This raises the possibility that both VAB-1 – EFN-1 and VAB-1 – EFN-4 interactions are required to shape the overall morphology of AIY neurons. Careful dissection of the *C. elegans* EphR, ephrin and HSPG promoter regions may define cell specific enhancers that could be used for rescue assays to better understand the complex relationship between these highly conserved pathways.

**Figure 10.**
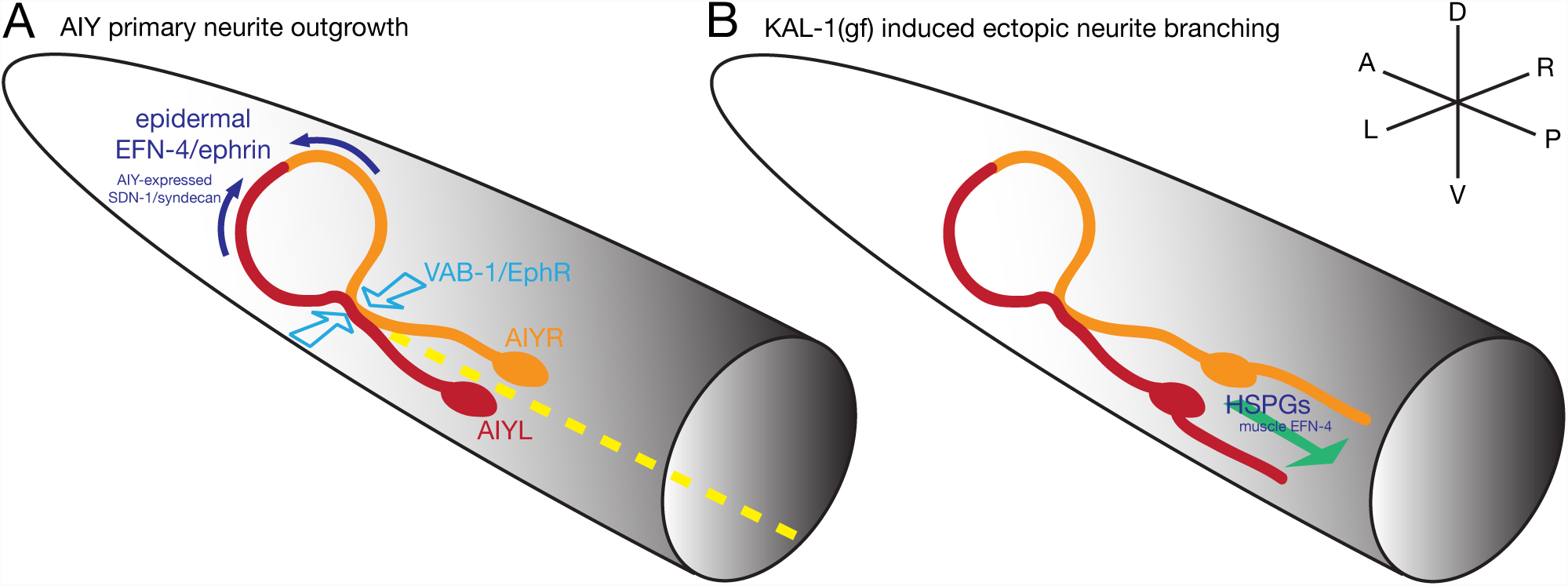
EFN-4 functions in concert with VAB-1 and HSPGs to shape AIY interneuron morphology in both wild type and *kal-1(gf)* contexts. Model summarizing the roles of EFN-4, SDN-1 and VAB-1 in promoting AIY interneuron primary neurite outgrowth. EFN-4, functioning non-autonomously from the epidermis, and SDN-1, functioning cell-autonomously in the AIY neurons, promotes extension of the AIY primary neurites toward the dorsal midline but have no obvious roles in axon guidance. VAB-1 has a minor role in nerve ring primary neurite extension but has a major role in driving AIYL /R contact in the ventral plexus. Whether VAB-1 is functioning cell autonomously in this process remains to be determined. The L1CAM LAD-2 may partially suppress EFN-4’s role in primary neurite outgrowth (not shown). Dashed yellow line illustrates the ventral midline. *kal-1(gf)* induced ectopic neurite branching is genetically distinct from primary neurite outgrowth. This phenotype is intrinsically cell autonomous, being driving from an integrated AIY-specific *kal-1* over-expression array. EFN-4 is required, in part, to promote secondary neurite branching where it functions non-cell autonomously in the body wall muscle. Ectopic branches can be found emanating from either the cell bodies or the ventral portion of the primary neurite (for clarity, only a cell body branch has been shown). HS plays a major role in *kal-1(gf)* induced ectopic branching, as loss of any of the HS modifying enzymes HST-2, HST-6, HSE-5, or SUL-1 strongly suppress this phenotype. The roles of HSPG core proteins in this phenotype are complex. Over-expression experiments indicate that GPN-1 and SDN-1 function as cell-autonomous regulators of *kal-1(gf)* branching. SDN-1 promotes ectopic branching whereas GPN-1 inhibits branch outgrowth. *sdn-1 gpn-1 lon-2* triple mutant data reveals a subtle role for LON-2 in promoting axon branching and highlights the overlapping and part-redundant roles for HSPGs in this process. Whether loss-of-function on one HSPG core protein leads to longer HS-branches on other core proteins is not known, but could provide a molecular explanation behind some of the core protein redundancy. Although our data indicates that no other HSPGs are involved, there may still be part-redundant roles for UNC-52/perlecan and CLE-1/collagen XVIII in this process.

## ACKNOWLEDGEMENTS

We thank Kathryn Hudson, Melissa Bentley, Philippe Mansour, Sebastian Chalfont, Lindsay Roe and Mike Branden for help with strain maintenance, Lihsia Chen for sharing reagents and data prior to publication, Hannes Bulow and Oliver Hobert for the original observation of *efn-4’s* role in *kal-1(gf)* axon branching, and Susan Smith and Lihsia Chen for their thoughtful comments on the manuscript. We also thank Erick Lee Purkhiser, R. Horton Heat and Rusty Hodge for their inspiration during this work. Some strains were provided by the CGC, which is funded by the National Institutes of Health (NIH) Office of Research Infrastructure Programs (P40 OD010440). Additional strains were provided by Shohei Mitani and the National Bioresource Project, Tokyo Women’s Medical University School of Medicine, Japan. M. L. H. was supported by a NIH K-INBRE research fellowship, a Knights Templar Eye Foundation Inc. Pediatric Ophthalmology Research Grant, and awards from Kennesaw State University’s Center for Excellence in Teaching and Learning and the Office of the Vice President for Research. This work was supported by NIH awards to J. L. M. (R15 GM080701), B. D. A. (P20 GM103638) and A. D. C. (R01 GM054657).

